# Transformations of the spatial activity manifold convey aversive information in CA3

**DOI:** 10.1101/2025.03.04.641439

**Authors:** Albert Miguel-López, Negar Nikbahkt, Carlos Wert-Carvajal, Lena Johanna Gschossmann, Martin Pofahl, Heinz Beck, Tatjana Tchumatchenko

## Abstract

Hippocampal circuits form cognitive maps representing spatial position and integrating contextual information, including affective cues, within episodic memory representations. We investigated how spatial and affective information combine in the population activity of CA3 axons by imaging intermediate-to-dorsal and dorsal-to-dorsal projections in mice navigating a linear track before, during, and after exposure to an aversive air puff stimulus. Our analyses reveal that both axonal populations maintain a robust, time-invariant spatial coding manifold across recordings, independent of affective context. Alterations to this common manifold encode the presence of the aversive stimulus without disrupting the spatial representation. Both axonal pathways encoded affective information with similar efficacy. This population-level encoding was distributed similarly across place and non-place cells. Our findings demonstrate that hippocampal CA3 axons integrate spatial and affective information within a common representational geometry while preserving the ability to extract each information type separately.

## Introduction

The hippocampus has been functionally associated with cognitive maps and episodic memory^1,2^. It integrates information streams, such as sensory or contextual cues^3^, into sequential representations. The interaction between spatial and affective information processing, particularly in natural behaviors like foraging or threat avoidance, represents a critical aspect of hippocampal function. While extensive research has characterized specialized cell types within the hippocampus – e.g., place cells^4,5^, object cells^6^, reward cells^7^ or goal direction cells^8^ – a comprehensive understanding of how salient features of the environment influence these neural representations remains incomplete. In particular, the interaction between spatial and affective information processing, especially in natural behaviors like foraging or threat avoidance, is a critical yet understudied aspect of hippocampal function.

We investigated how aversive information alters spatial hippocampal codes, in a navigation task, by analyzing long-range CA3 pyramidal cell projections. CA3 specifically was selected for its rapid spatial information acquisition^9^, robust spatial tuning^10^, and because in particular ventral CA3 neurons are known to process aversive information ^11,12^. We monitored the activity of individual CA3 axons projecting from the intermediate to the dorsal hippocampus, as well as commissural CA3 axons originating in the contralateral dorsal hippocampus, using 2-photon imaging in mice. The animals were running on a linear track containing tactile cues and could earn a reward after each lap. An aversive stimulus in the form of an air puff was introduced and later removed. We focused on analyzing axonal population activity to understand how spatial representations were conserved across time and how they integrated contextual affective information after the introduction and subsequent removal of the aversive cue.

We observed a remarkable consistency in spatial representations across sessions, indicating a robust organizational structure and common task abstraction. Rather than maintaining separate representations, spatial and affective information were jointly embedded in a shared neural manifold. We observed that intermediate-dorsal axons had greater plasticity in response to aversive conditioning compared to their dorsal recurrent counterparts, although that did not result in a better encoding of aversive information. Notably, removing the aversive stimulus resulted in a distinct post-stimulus state rather than returning to pre-stimulus activity, suggesting an affective memory effect. Finally, we showed that affective information was embedded across cell types and dimensions, with non-place cells contributing as much information as place cells to contextual decoding.

Overall, our results indicate that affective information is directly embedded within the hippocampal state-space, and is able to deform its existing manifold creating an integrated representational space that keeps its spatial information consistent across time. Our work provides crucial insights into the mechanisms by which the hippocampus creates comprehensive cognitive schemas that incorporate both spatial and affective components of experience.

## Results

To analyze the integration of spatial and affective information in CA3 we recorded intermediate-to-dorsal (ID) and dorsal-to-dorsal (DD) axonal activity in mice as the animals ran along a 1500 mm long looping linear belt (Fig. 1A, see Methods). A reward was given at the end of each round (which we define as a trial) along the belt to motivate the animal to complete laps, with tactile cues allowing the animal to differentiate each third of the belt and estimate its location correctly. In baseline sessions (*B*_1*−*3_), taking place on the first two days, the animals ran across the belt without any external intervention except for the reward. On the third and fourth training days, an air puff (AP) was presented in all trials whenever the animal reached the position *x* = 500*mm* (training sessions, *T*_1*−*4_) and was subsequently removed on day 5 (probe sessions, *P*_1*−*2_). Population-wide calcium sensor signal was recorded and transformed to dF/F (Fig 1C) from ID axons in 4 mice (M0-M3), and from DD axons in a separate group of 4 mice (M4-M8), subject to the same behavioral protocol^12^.

**Figure 1:**
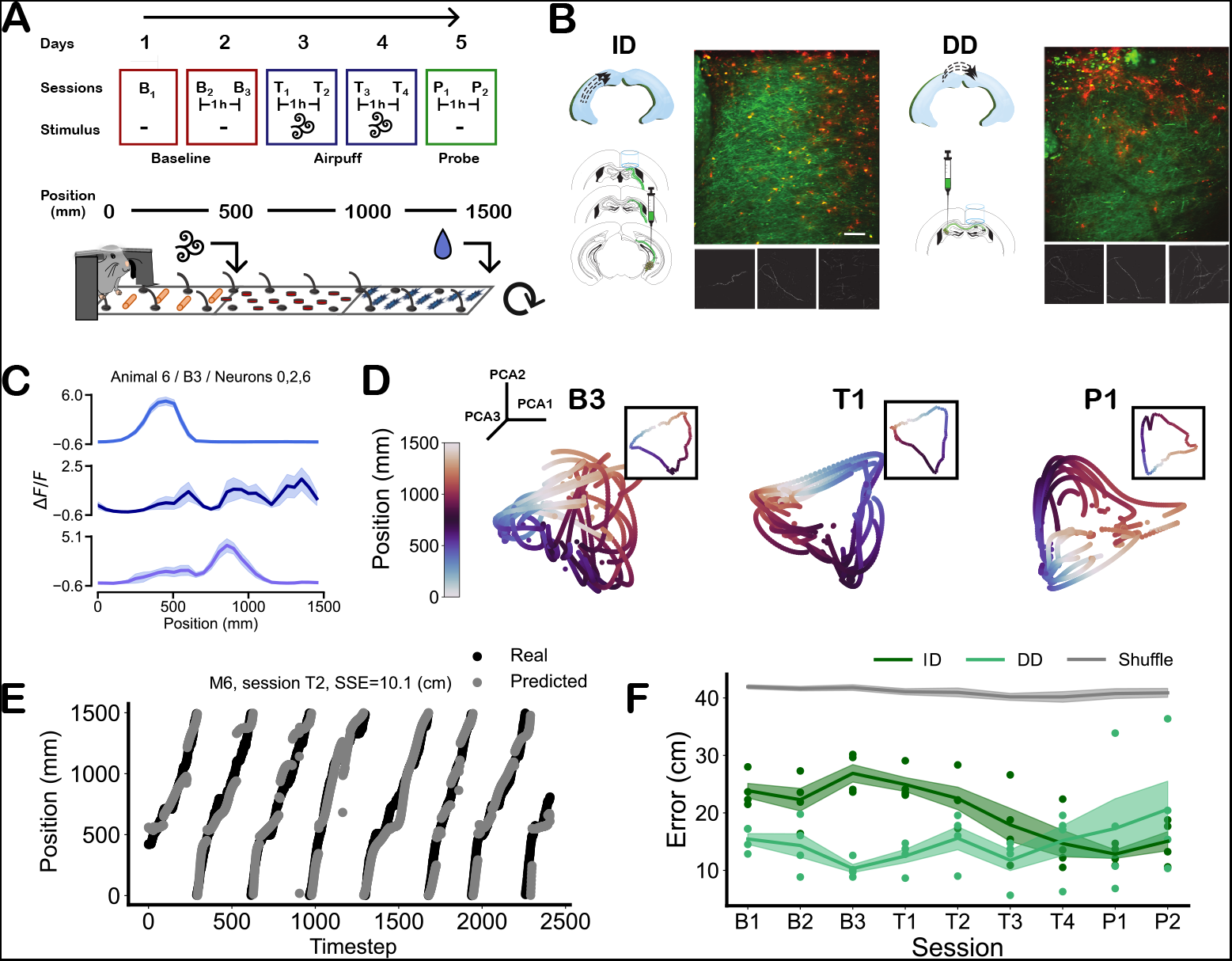
Representation of belt position in ID and DD axons remains robust as manifold position and orientation varies across sessions. **A**, Sketch of the task design. The top panel represents the behavior protocol. Each box represents a day of recording, and sessions in the same box are separated by only 1h time interval. T (training) sessions include an aversive stimulus, not present in B (baseline) and P (probe) sessions. The bottom panel shows the behavioral apparatus design, with a headfixed mouse running on a circular belt. A liquid reward was given at the end of each lap. The aversive stimulus only appears in T sessions at the 500 mm mark. **B**, Diagrams of coronal sections illustrating the ID and DD injection sites for calcium indicators in the mouse brain (left panels), together with examples of two-photon imaging field of views (top right panels), the black lower squares depict example axonal components extracted from the recordings. **C**, Δ*F/F* averaged over trials for an example animal and session, three neurons shown. Shaded areas show the standard error of the mean (SEM) of the data. **D**, Three-dimensional plots of the latent space for an example animal and three representative sessions. Color represents the position on the belt. Insets show the trial-averaged data for that session. **E**, Position prediction on an example session. **F**, Prediction performance averaged over mice, for each session and axon type (ID or DD). Scatter dots are the averaged prediction errors for each mouse and session. The gray line is the average prediction error after shuffling the position information (10 repeats per session). Shaded areas show the SEM of the data. ID and DD prediction errors were significantly different than shuffle (t-test, *p <* 0.05, mice and sessions pooled together), as well significantly different from each other.

We characterized the latent space of each recording using Principal Component Analysis (PCA), the first three components of which reflected the periodic nature of the belt (Fig. 1). While the exact shape and coordinates of these manifolds changed from session to session, the overall periodic structure was robust across all sessions. The latent space we found was low-dimensional, with e.g. less than half of the original dimensions necessary to explain 80% of the variance (Fig. S1B-D). To quantify this activity manifold, we trained a decoder to predict the position of the animal from the latent axonal activity, taking enough principal components (PCs) to explain 90% of the variance for each session. We were able to predict position accurately (Fig 1E), with the prediction error being significantly different from shuffle controls for all sessions and axonal types (Fig 1F). Interestingly, ID axons had higher errors during the baseline sessions, which lowered to the level of DD errors after introducing the aversive cue during T sessions. This suggests that ID axons improved their spatial encoding as a reaction to the aversive stimulus, while DD kept a more stable spatial representation over sessions, matching our previous results at the single cell level^12^.

### Common spatial manifolds exist across sessions

Even though neuronal latent structures appear similar over sessions, a decoder trained on the position of an animal on one session will generally perform poorly when predicting on a different one (Fig. 2A middle, and “unaligned” in Fig. 2D). This approximately twofold increase in the error (“unaligned” in Fig. 2F) could be explained with an invariant spatial map that is drifting, rotating, or deforming across time in the axonal activity space. Only some axons are shared from session to session, and those may have their stimulus-dependent responses drift over time or even remap. One could understand the latent space of each session as the projection of an underlying common manifold^13,14^ onto the recorded activity of each day.

**Figure 2:**
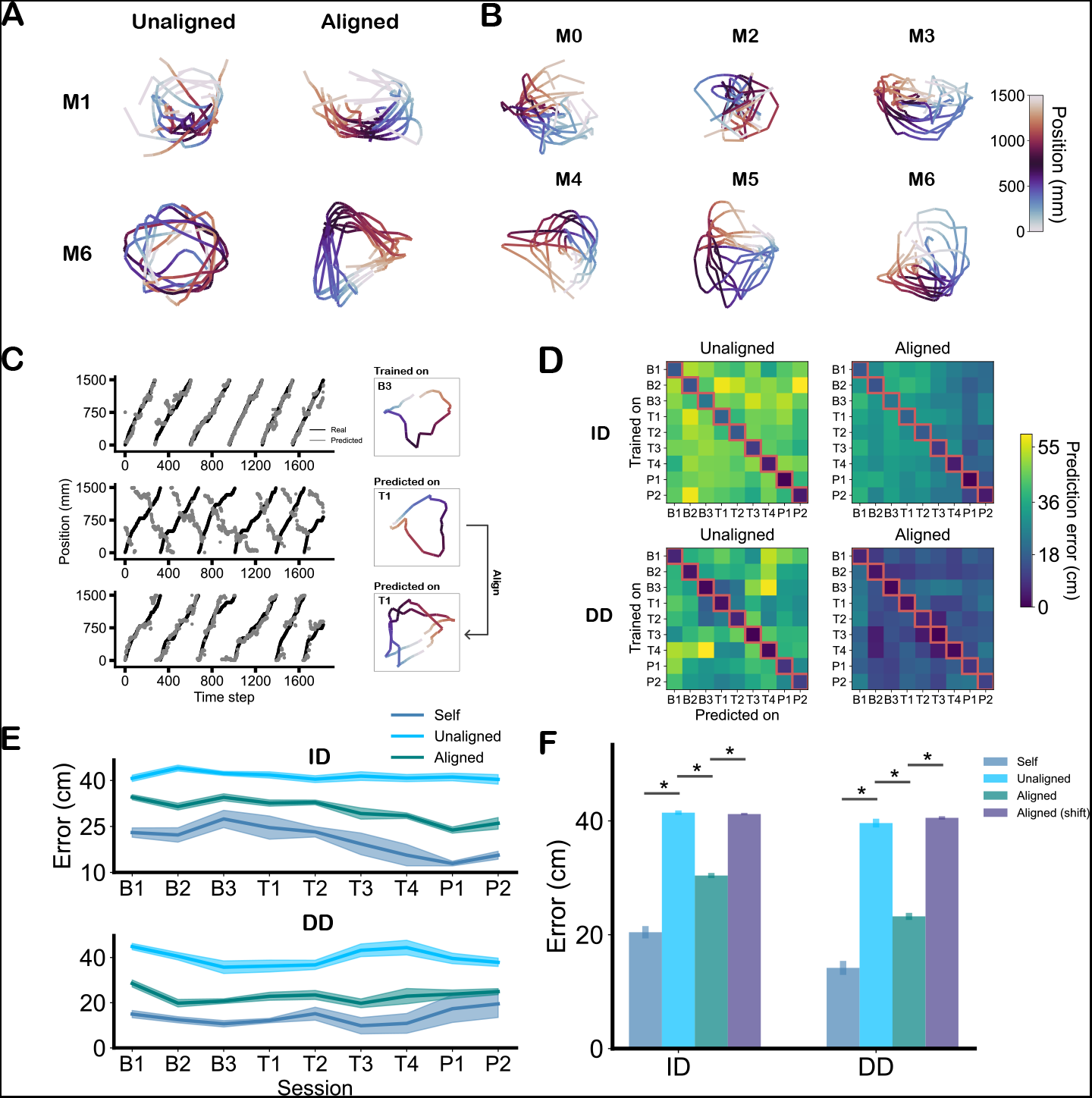
Revealing a common spatial manifold across sessions in axonal population activity. **A**, Spatial manifolds in ID axons (top) and DD axons (row) can be aligned across task types indicating robust spatial representation space that is independent of task-context. The first three dimensions of session-averaged PCA, before and after alignment, are shown. **B**, Aligning the population activity across all sessions within individual mice (mice IDs M0, M2, M3,M4, M5, M7) reveals manifold similarities. **C, left**, Dynamic manifolds encoding the spatial position of the animal (color code) across different session types (baseline session *B*_2_ vs air-puff session T1) can be aligned via CCA (mouse M6 shown as example). **C, right** Ground truth position (black) vs prediction (grey) for cross-validated *B*_3_ session (top), for a decoder trained on *B*_3_ predicting on *T*_1_ (middle), and predicting on aligned *T*_1_ data. Right column shows the session-averaged PCAs. **D**, Position prediction improves substantially before (left column) vs after (right column) alignment. The first row is the average across ID mice (N=4), and the second is the average across DD mice (N=4). Diagonal squares (red border) correspond to cross-validated self-error, while the non-diagonal squares correspond to cross-session prediction. **E**, Prediction error across sessions for each axonal type. Shaded area represents the standard error of the mean of the data. **F**, Animal averaged errors in ID vs. DD populations. Asterisks indicate a significant difference (t-test, *p <* 0.05, unpaired for self to unaligned, paired for the others). Vertical lines show the standard error of the mean of the data.

To test this hypothesis, we aligned the activity manifolds in the principal component space across all sessions using multiset Canonical Component Analysis (mCCA), which maximizes their cross-correlations through linear transformations (see Methods). Before alignment, activity manifolds displayed similar periodic structures but in different shapes and rotations (Fig. 2A, left column). These differences were reduced through the alignment procedure, as seen in the right column of Fig. 2A, and 2B. To quantify the strength of the alignment, we assessed spatial encoding stability via cross-session decoder generalization, following the process shown in Fig. 2C. To avoid favoring any particular session, we trained and tested a decoder for every possible session pair (Fig. 2D). We observed a clear reduction in the cross-session prediction error after alignment (non-diagonal cells in Fig. 2D, and Fig. 2F). This improvement persisted for the entirety of the experiment (Fig. 2E), even for ID axons during the initial sessions, which had a larger within-session error than the rest (Fig. 1F). Overall, the error in aligned sessions was significantly lower than in unaligned ones and much closer to the error obtained when predicting the same session (Fig. 2F), for both ID and DD axons. To eliminate the possibility that mCCA was able to align datasets independently of their underlying structure, we repeated the process on data that had a random temporal shift applied on each session so that the latent structure became uncorrelated with position but keeping its temporal correlations intact (Fig S2). Prediction error was comparable to shuffle in this case (Fig. 2F, aligned shift), indicating that a shared latent structure with a consistent relationship to task variables is needed for mCCA to work. Overall, the success of the alignment procedure demonstrates that a common space manifold was present across all sessions for both axonal types and that a simple linear transformation is enough to recover it.

### Affective information is encoded within the spatial activity manifold

Next, we wanted to understand how the appearance of a novel aversive stimulus (the air puff, or AP) could modify the underlying representational manifold. Since position decoders worked well across all sessions after alignment, we considered the possibility that any contextual changes would happen trial-to-trial rather than timestep-to-timestep. To investigate this hypothesis, we applied Tensor Component Analysis (TCA) to our data^15^, a linear technique that decomposes neuronal recordings into three sets of “factors” or components: neuron-wise, time-wise, and trial-wise. For each mouse, we re-arrange our data into a third-order tensor *T* where the first axis contains the aligned PCs from the previous alignment step, the second axis contains the position-normalized bins for each trial, and the third axis concatenates all trials from baseline to probe sessions (Fig. 3A, left). We concatenated sessions using the aligned dimensions because we are interested in observing changes in the common latent space across time and trials. TCA performs dimensionality reduction on the tensor as a whole, decomposing each element of *T* into the sum of three sets of factors (Fig. 3A, see Methods). Intuitively, the process can be understood as performing a PC decomposition with an added third source of variability – i.e., trials – and removing the condition of orthogonality between components.

**Figure 3:**
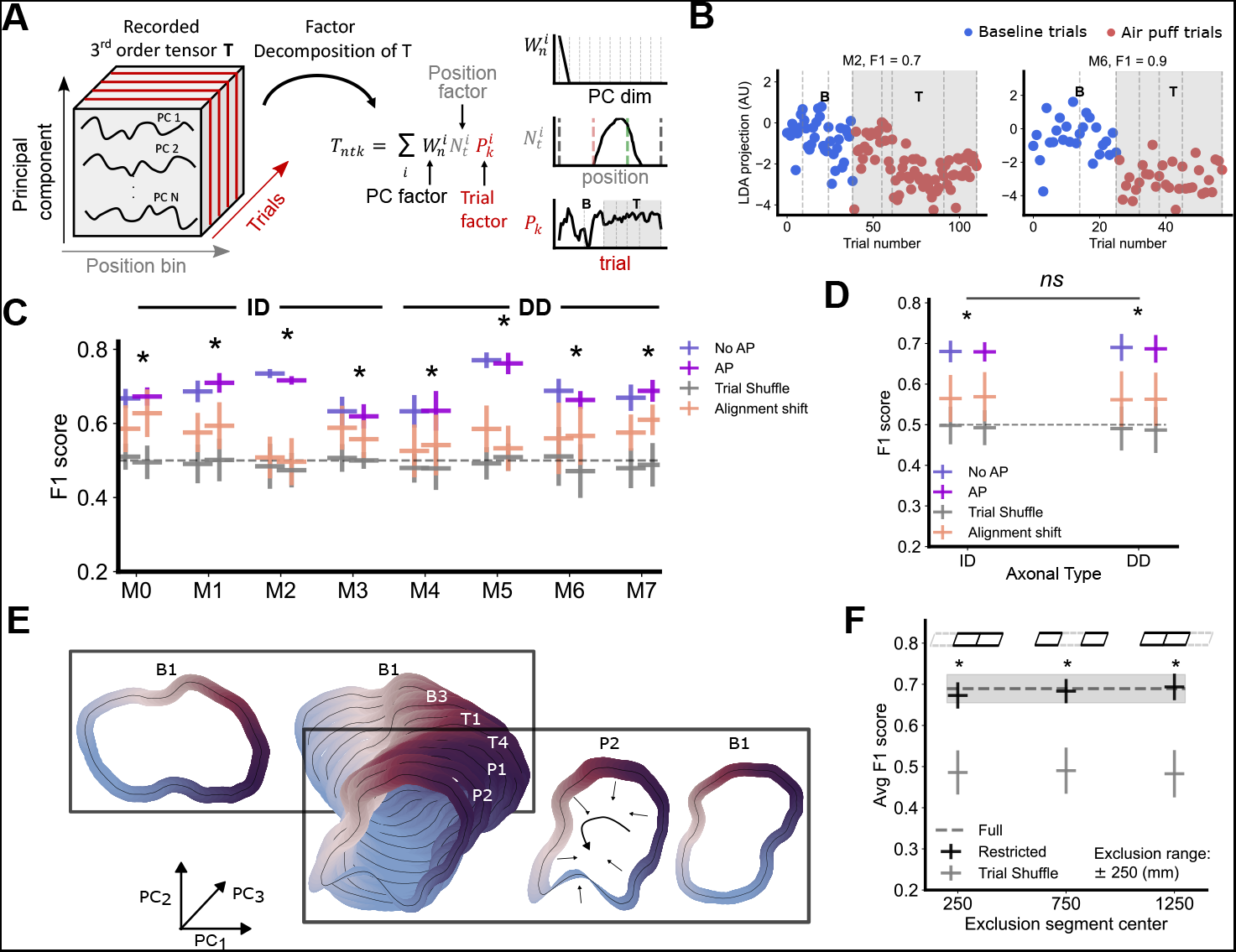
Decomposition of the neural activity tensor identifies dimensions encoding aversive information. **A** The recorded 3D tensor (A, left) can be decomposed into trial, position and neural PC factors (A, middle). (A, right) shows one representative TCA factor for each decomposition. **B** LDA projections of trial factors onto a 1-dimensional space. Each dot corresponds to the projection of one trial factor. Session delimiters are marked with dashed lines, air puff sessions have a gray background. Left panel shows mouse M2 (ID axons) and right panel mouse M6 (DD axons). F1 scores are shown on top. **C** F1 scores for each mouse and both classes, averaged over TCA repetitions. Vertical lines show the standard deviation (SD) of the data. Mice whose F1 scores are significantly (t-test, *p <* 0.05) above trial shuffle (gray) and alignment shift (brown) controls for both classes are marked with an asterisk. are well above the significance threshold. Dashed line indicates the expected F1 score of a completely random guesser. **D** Mouse-averaged F1 scores by axon type. Vertical lines show the SD of the data. Asterisks indicate that both classes were above trial shuffle (gray) and alignment shift (brown) controls, as in the previous panel. The difference between ID and DD was non-significant. **E** Representation of the latent space deformation under contextual changes. Each periodic structure represents the trial-average PCA of a session, with colors indicating position along the periodic belt. As context changes, the latent manifold is modified in ways that keep its space encoding mostly intact. **F** Air puff decoding performance when the activity corresponding to certain parts of the belt is excluded. Drawings (top) indicate which third is excluded. The F1 score of the whole dataset (dashed line), as well as the F1 score of the restricted dataset with randomly shuffled labels (gray), are added as a comparison. Vertical lines show the SD of the data. Asterisks denote significance w.r.t. shuffle.

Figure 3A right shows the first component of each TCA decomposition for a representative session (more components are shown in S3A). The PC factor shows the weights for each aligned PC dimension, exhibiting a highly selective distribution across the discrete principal components. (with some dimensions being fully active while others are not). This indicates that TCA is picking up on the dimensions already discovered by PCA or CCA and explaining most of the variance. The position component has a place cell-like shape, indicating that TCA is merging place field information, which is likely coming from the PCs. The third dimension exhibits fluctuations across the trial space, indicating how the amplitude of this activity pattern changes across trials. A geometric intuition for these varying trial factors is given in Fig. 3E. As components gain and lose strength over trials and sessions, the overall manifold is deformed along selected dimensions, keeping positional information intact (illustrated by the small arrows *P*_2_ vs *B*_1_ and the preserved periodic shape).

To quantify this variability, we focused on evaluating fluctuations of the trial components across session types. Initially, we solely considered changes accounting for the learning of aversive stimuli, comparing baseline vs training sessions. We employed Linear Discriminant Analysis (LDA) to decode air puff presence from the trial components using leave-one-out cross-validation (Fig. 3B) and evaluated it using the F1 scores for each class. We used scores for randomly shuffled labels as a baseline for comparison (“trial shuffle” in 3C and D). Additionally, we considered the possibility that the decoder was simply picking out session-to-session variability independent of air puff presence. Such variability might arise through a flawed alignment process, or changes due to the passage of time such as drift, or the animal becoming more accustomed to the belt environment. To control for this, we repeated the analysis using sessions that were each independently shifted by a random interval, which CCA failed to align successfully (Fig. 2F); hence, it must be mismatched when concatenated along their PC axis. Air puff decodability was significantly above both control types for all mice (Fig.3C, raw accuracy metrics in Fig. S3B and C), indicating that the common manifold changed due to the introduction of an aversive stimulus. Notably, we found that air puff presence was decodable in both ID and DD axonal recordings, showing no meaningful differences (Fig.3D), which indicates that both axonal types react to aversive information similarly. This appears to challenge the idea that ID axons encode more affective information than DD.

To rule out that air puff decoding was caused by behavioral correlates – e.g.,the animal slowing down in the air puff area – we re-computed the trial factors and repeated the analysis using only activity from particular sections of the belt (Fig. S4A). Excluding one-third of the belt did not significantly alter the results, even when the excluded section included the AP or pre-AP belt zones where the animal slows down during T sessions (Fig.3F and Fig. S4C). Only one mouse (M2 in Fig. S4B) appeared to show a significant reduction in this case. Interestingly, the results were largely interchangeable between different belt sections, even when two-thirds of them were excluded (Fig. S4D). This suggests that aversive information is distributed along the entire latent manifold regardless of position. It also shows that the deformations schematized in Fig. 3A are pervasive along the belt, not just in locations where the animal modifies its behavior.

### Probe sessions constitute a distinct post-aversive state

Next, we extended this analysis to include probe trials. After removing the aversive stimulus, animal behavior became more variable, with stopping behavior spreading out in a wider space interval^12^. This made us question if axonal activity during probe trials significantly differed from baseline and air puff trials. When trying to decode air puff presence by grouping B and P sessions together against T (BP-T in Fig. 4A), performance was similar to our previous analysis comparing only B against T. But this was also true when trying to differentiate B from P trials, which lacked the presence of an air puff (B-P in Fig. 3A). This indicates that axonal representations in the post-AP context become distinct from the pre-AP context, rather than returning to the baseline after removing the aversive stimulus.

**Figure 4:**
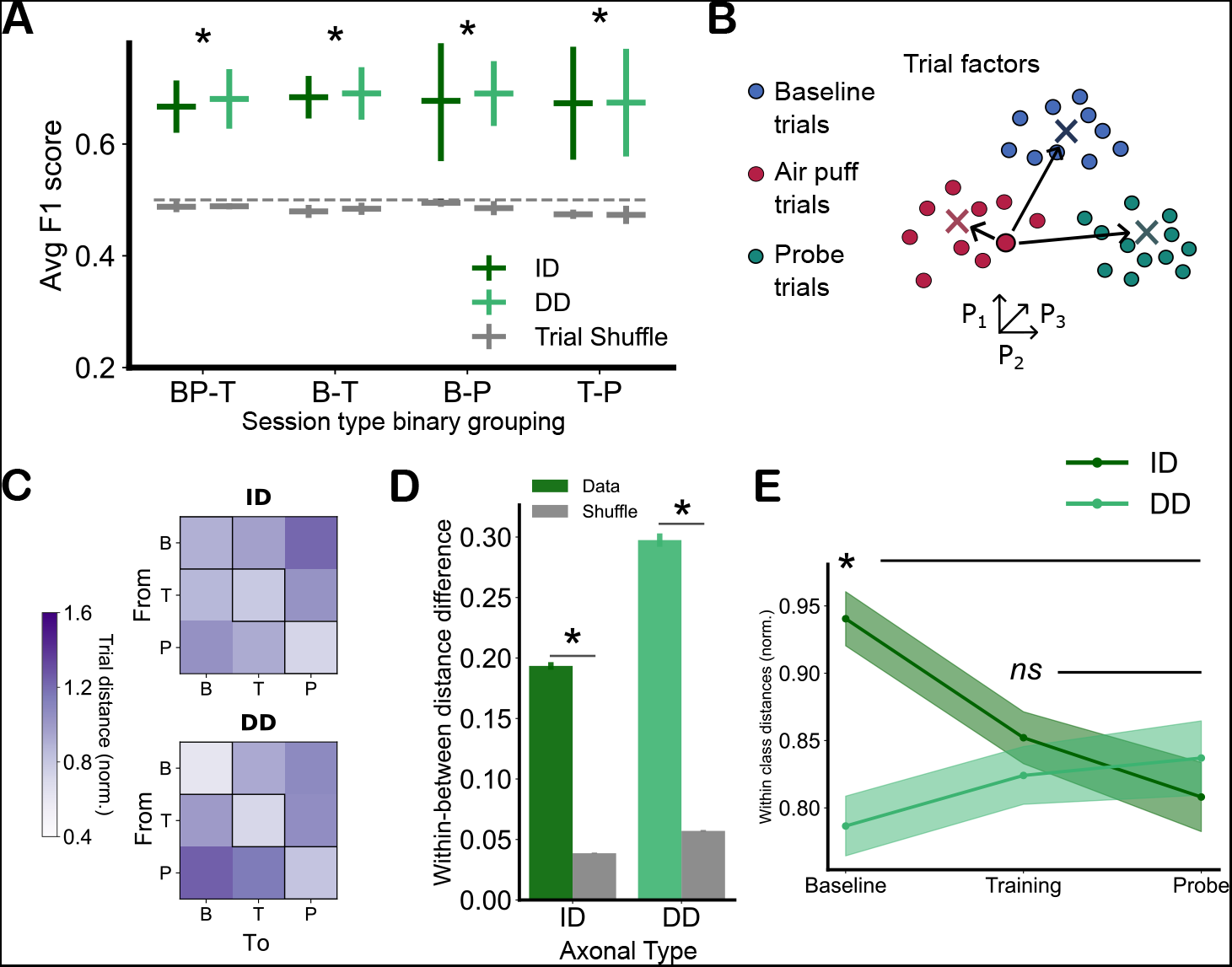
Aversive information is distributed across dimensions and cells and is encoded differently from positional information. **A**, F1 scores for different binary comparisons of the three session types, and for each axonal type. “B” are baseline sessions, “T” are air puff sessions, and “P” are probe sessions. The x-axis labels indicate the groups of sessions compared against each other (e.g. in BP-T we distinguish trials belonging to either B or P sessions from T sessions). An * indicates that both axon types for that session grouping were significantly (t-test, *p <* 0.05) above shuffle (in gray). Dashed line indicates the difference expected from random draws. Vertical lines show the standard deviation (SD) of the data. **B**, Computing distances in trial factor space. Each trial of a session is represented by a single point whose xyzcomponents are amplitudes of the corresponding trial dimensions. For visualization, we show three trial factor dimensions (x, y, and z axes) and color the activity vectors in blue, red, and green if they correspond to a baseline, trial, or probe sessions respectively. Crosses represent the centroids of each cluster. A single trial is highlighted as an example, with solid black lines indicating the distances from the trial to the centroids. **C**, Colormap of the average normalized distance from a trial to each centroid, by axonal type. The diagonal elements, highlighted by a black outline, show the average within-cluster distance. **D**, Average difference between the “within” and “between” normalized distances for each axonal type, for both the original data and the label-shuffled data. Significance (t-test, *p <* 0.05) was computed with respect to the results from shuffled label data. Vertical lines show the standard error of the mean of the data. **E**, Within class normalized distances across session types for ID and DD axons. Asterisks indicate significance (t-test, *p <* 0.05) between ID and DD differences relative to the Probe distances. Shaded areas represent the SD of the data.

Motivated by this finding, we wanted to understand how the different session types were distributed in trial factor space. We hypothesized that each session type would form its own distinct cluster (Fig. 4B) and quantified it by measuring the distance between trials and the centers of mass of each session type (Fig. 4C). We found that, on average, within-cluster distances were smaller than between-cluster for all session types when compared to shuffled session type labels (Fig. 4D, and Fig. S5F), supporting the idea that the B and P latent manifolds are distinct. This was corroborated when we repeated the analysis grouping B and P trials together (air puff or AP class) against T (no air puff class or NoAP), finding that the AP class was more spread out than the NoAP (Fig. S5I), implying that we were grouping distinct clusters together. Additionally, separability was better in DD axons compared to ID (Fig. 4D), which was likely caused by a higher within-class spread before the air puff introduction (Fig. 4E). This result aligns with the position decoding results from Fig. 1F, suggesting ID axonal activity stabilizes after presenting an aversive stimulus into the environment. Overall, these results indicate that removing the aversive cue caused changes in the latent manifold that consolidated into a new, distinct contextual encoding, different from the context before the air puff was introduced.

### Aversive information is evenly distributed across dimensions and cell types

Finally, we wanted to know how aversive information was encoded across dimensions and axons. Figure 5A shows that the average decodability of the aversive cue increased progressively with the number of TCA dimensions, indicating that aversive information was found beyond just the initial PCs. To understand exactly how affective information is distributed, particularly in comparison to spatial information, we quantified the contribution of each latent dimension or each axon to the decoding process with its decoding weight for either position or air puff decoding (see Methods). Dimensionality-wise, initial PCA dimensions contributed the most to position prediction, while air puff information was more evenly distributed along dimensions (Fig. 5B), indicating that spatial coding dominates neuronal activity patterns, as they occupy the PCs with larger explained variances. When measuring the contributions of place and non-place cells (see Methods for classification details), we found, as expected, that place cells had a clear positive contribution to spatial decoding (Fig. 5C). But this was not true for air puff decoding, with both cell types contributing a similar amount of information, with even a small but significant difference in favor of non-place cells. Finally, we found very weak correlation (r=-0.067) between the air puff and position weights of a given axon (Fig. 5D), suggesting that aversive information is found in cells independently of their role in position encoding. This is consistent with previous reports showing that individual place and non-place cells contribute significantly to place as well as environmental encoding^16^.

**Figure 5:**
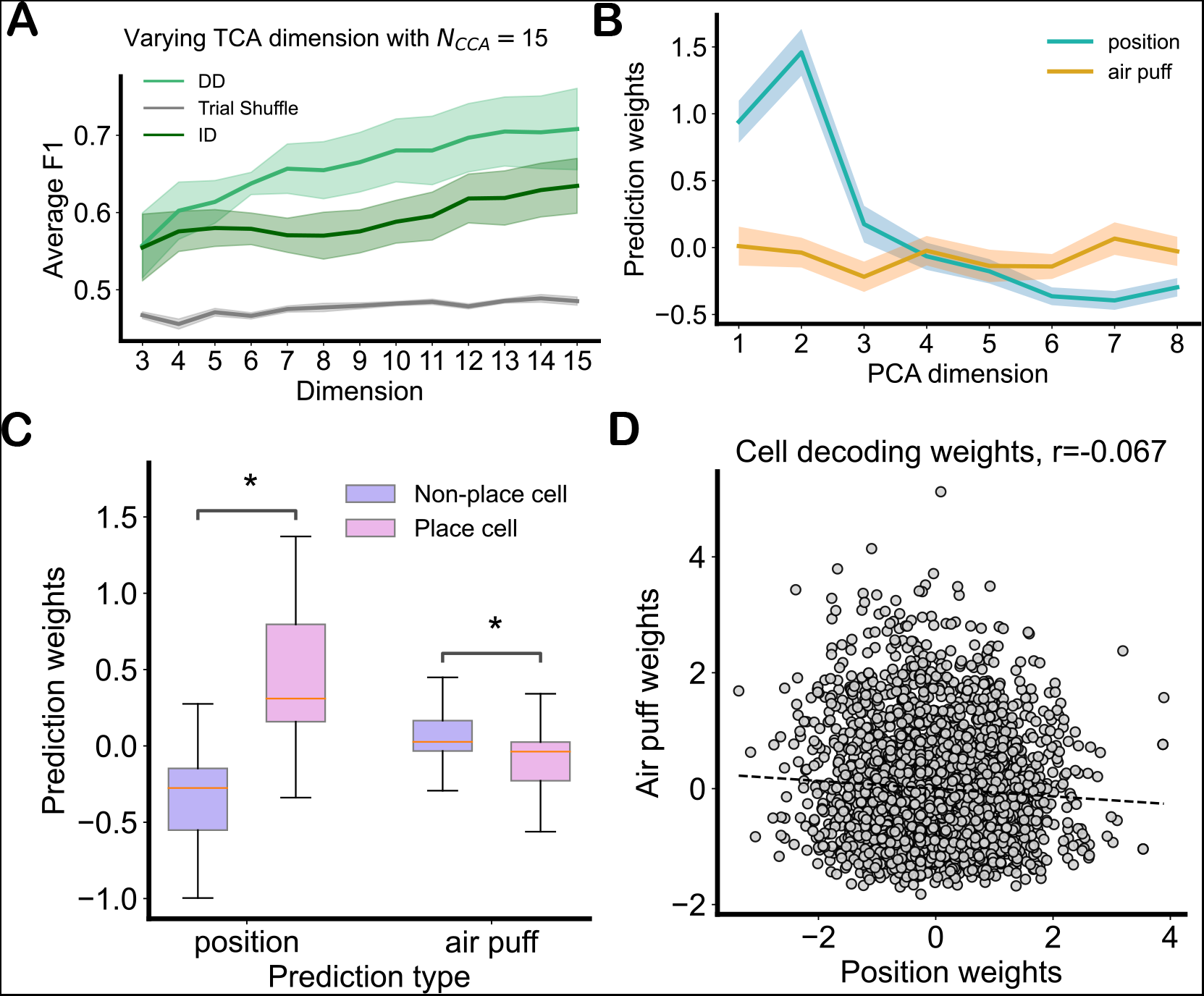
Air puff effect is robust and distributed across cells and dimensions. **A**, F1 scores plotted against TCA dimensionality, while keeping CCA dimensionality fixed. Shaded area shows the standard error of the mean. **B**, Average prediction weights (see Methods for details) for position and air puff decoding by principal component dimensions. Shaded area shows the standard error of the mean. **C**, Boxplot for the position and air puff decoding weights by cell type. Asterisks denote significance between cell types (t-test, *p <* 0.0005). The colored line within the box represents the median, the box represents the inter-quartile range (IQR), which includes quartiles 1 to 3, and the whiskers extend 1.5 times outside the IQR. **D**, Cell weights for air puff decoding plotted against weights for position decoding. Each dot corresponds to a cell. The dotted line shows the linear fit, with a p-value of 0.0007.

To summarize, we have shown that aversive information is similarly distributed between place and non-place cells independently of their position information, and that non-place cells have a key role in encoding affective information.

## Discussion

Integrating salient information from the environment into spatial maps is a fundamental cognitive function animals accomplish effortlessly. Yet it remains unclear how this happens at the neuronal level. While changes in single-cell selectivity induced by affective information or task context have been observed across different cortex^17^, how the population-wide activity manifolds combine these different information types remains an open question. Previous research has shown that when spatial and non-affective context variables are simultaneously present during a task, they jointly map into a shared geometric representation^18^. Following the idea that hippocampal circuits construct structured maps using the substrate of neural activity manifolds ^19–22^, it is plausible that existing spatial maps are transformed to accommodate newly arriving information^23^. The possibility of a joint encoding of spatial and affective information within a single manifold is further supported by observations that many cell types in the hippocampus can change their firing patterns or acquire novel receptive fields in new task and environmental contexts ^12,24^.

Here, we have shown that the spatial map generated by a population of CA3 axons is preserved across different affective states despite receptive field remapping of individual cells within the population^12,16^. Notably, we show the existence of a latent activity manifold whose positional representation remained stable across days, even after contextual changes caused by the introduction and subsequent removal of an aversive stimulus in the form of an air puff. The latent representation in the principal component space obtained on each session showed a consistent periodic structure, which reflected the nature of the belt and accurately encoded the position of the animal. However, the specific shape of this spatial representation changed across sessions, such that a decoder trained on one recording would fail when predicting on another. This could be caused by a number of factors such as recording different cell populations, remapping, or representational drift. Importantly, these differences largely disappeared after aligning the recordings. This was also true for ID axonal populations, whose spatial representation improved over time. This suggests that the introduction of an aversive stimulus caused the recruitment of more neurons to strengthen this shared manifold.

Next, we found that the presence of the air puff could be predicted from the activity changes in the aligned manifold at the single trial level. This indicated that affective information was encoded in the neuronal activity. We hypothesized that the aversive cue had a “deformation” effect on the latent activity manifold, such that contextual changes would alter its general ring-like structural while positional encodings remained unaffected. This was observed regardless of which section of the belt we analyzed, suggesting that contextual information was distributed across the entire manifold, and that it was uncorrelated with behavioral differences such as the animal stopping at the air puff location. Another interesting result was that, even though ID axons are thought to carry affective information into the dorsal region, and in our own data showed a stronger reaction to the introduction of an aversive stimulus, both DD and ID axonal populations had similar air puff decodability. It is possible that both regions encode affective information, but this is only visible at the population level, rather than within single cells. Another possibility is that this information is first transmitted to the dorsal region through intermediate-dorsal projections, and then quickly integrated in the dorsal-dorsal projections. Causal studies such as lesion experiments would be necessary to understand this phenomenon.

Further analysis of the probe sessions revealed that neuronal activity following the removal of the aversive cue was best characterized as a distinct neural state, different from the baseline sessions which also lacked air puff presence. This suggests that not just affective, but also memory-affective information (such as the knowledge that a negative stimulus was present in the past) is represented in this common latent manifold. This was reflected in the behavior of the animal, showing more spread out stopping locations and variable stopping frequency ^12^.

Finally, we found that contextual information was distributed evenly along dimensions and cell types. In contrast, spatial information was located mainly along the principal components that explained the most variance, as well as in place cells compared to nonplace cells. This result suggests that contextual changes affect the whole manifold beyond the spatial map, and that selectivity among hippocampal axons is highly mixed. The fact that non-place cells contributed at least as much as place cells to contextual decoding suggests that the importance of non-place cells in hippocampal neuronal maps could be understated and that they might be key to the integration of contextual information in the hippocampus. Further research should study how other contextual information may be integrated within common neuronal manifolds and examine how affective information flows from region to region.

In summary, our results indicate that population-wide spatial representations in the hippocampus might be conserved across time and contexts and that affective information is embedded into the state-space through variations in its underlying population dynamics.

## Acknowledgements

We acknowledge the support by the Institute of Experimental Epileptology and Cognition Research at the Medical Faculty of the University of Bonn, and Joachim Herz Foundation (C.W-C), DFG graduate school iMol (A-M.-L.). This research was funded by the Deutsche Forschungsgemeinschaft (DFG, German Research Foundation) – Project-ID 227953431 – SFB 1089 (H.B. and T.T.). We thank Juan Alvero Gallego for fruitful discussions and method suggestions during the early stages of the project.

## Methods

### Data acquisition

#### Behavioral Paradigm

The behavioral paradigm used in this project was a modified version of context-dependent fear conditioning as described in^12^, adapted to our two-photon setup and head-fixed condition. After a week of recovery following the virus injection and head-fixation implantation, the animals were habituated to the horizontal treadmill and the head-fixed condition running on the belt according to the following protocol. On the first day after recovery, mice were allowed to freely explore the treadmill. During this day, mice were placed two or three times during the active phase on the apparatus to sniff around and freely explore the environment and the belt, each time for a maximum of 10 minutes. On consecutive days, habituation to the head fixation was carried out. Mice were head fixed on the setup on each day for 2×10 minutes, until all signs of anxiety subsided and mice ran freely on the linear treadmill, with a plain belt. Such habituation days were repeated until the animals reached a steady performance and completed a minimum of 10 rounds in 20 min, up to two weeks after virus injection/head fixation implantation. Simultaneously with the habituation session, starting one week after the surgery, ad libitum food was discontinued and a scheduled food diet according to our animal-grant (AZ 84-02.04. 2014.A254) was administered. The protocol was as follows: The animals received a measured amount of food on the first two days of the food scheduling protocol. On the evening of the second day the remaining food was weighed in order to define the consumed amount of food. This value was divided by 2 and the corresponding amount of 1 gr food pellets (Dustless Precision Pellets®Rodent, Grain-Based, Bio-Serve) were placed onto the cage floor for consumption. The weight of mice and the consumed amount of food was supervised every day until they reached to almost 85% of their initial body mass. Additional care was taken in order to make this adjustment according to the general health and behavior status of each animal. During the food scheduling period, a heat-pad was placed underneath the cage to keep mice warm. During the habituation and behavioral sessions on the linear track, the animals received liquid food rewards at the end of each lap (diluted Butler Supplical Pet Multi-vitamin). Once the habituation phase was completed, cranial window implantation was carried out, followed by a recovery period. After recovery, we initiated the baseline and aversive learning protocol. On each of the first two days of the protocol, two 20-minute baseline sessions were carried out (session start of the second session 60-90 minutes after the first) on a new, cue-enriched linear track. The first session on the first day was discarded as the mice were confronted with the cue-enriched linear track for the first time. The last three baseline sessions before the inception of training sessions were consistently termed *B*_1_ − *B*_3_. During these sessions, all animals ran at least 8 laps during each 20 min session with an average of 21 laps per session. On the next days of the protocol, twice daily 20-minute training sessions were carried out, also 60-90 minutes apart. Two daily training sessions proved ideal to allow mice to learn the location of the relatively mild aversive stimulation. During training sessions, animals received a puff of compressed air each time they passed the border of the first and the second zones on the belt (at 50 cm according to the convention in all figures). Training sessions were repeated for as many days as needed until the animals showed apparent slowing down or stopping behavior close to the air puff location. Following the last training session, on the next day, a probe session was carried out. During the probe session, no air puff was applied to test if slowing at the previous air puff location persists, indicating memory formation. Thus, equal, 24 hour time intervals were present between sessions *B*_1_-*B*_2_, *B*_3_-*T*_1_, and *T*_4_-*P*_2_, these equal time intervals were used to quantitatively compare the dynamics of place coding over time. Speed on the linear track was analyzed with custom-written code in Python. First, we binned the speeds for every 1 cm on the belt, and added the binned data to a matrix with a dimension of “number of rounds x number of bins”.

#### Craniotomy and hippocampal window implantation

45 min before the start of the surgery dexamethasone was i.p. injected (0.1mg/20 g body weight) in order to inhibit brain inflammation during surgery. The animal was anesthetized inside an induction chamber with 3-4% isoflurane mixed with an oxygen/air mixture (25/75%) and was then transferred to a stereotactic frame. An isoflurane mask was put on the snout and the head was fixed using head-bars. The anesthesia was maintained with a reduced isoflurane dose of 1-2% at a gas flow of about 0.5 l/minute. Eyes were covered with eye-ointment (Bepanthen, Bayer) to prevent drying, and body temperature was maintained at 37°C using a regulated heating plate (TCAT-2LV, Physitemp). The skull in the middle of the head-ring was thoroughly cleaned using a Q-tip soaked in 70% ethanol. A Ø 3 mm craniotomy was made using a dental drill. The circular piece of skull was gently removed and any remaining bones around the edges were smoothed and drilled away. The cortex was then carefully aspirated with a blunt 27-gauge needle attached to a suction system until the alveus of the hippocampus became visible. When required, irrigation with dPBS was carried out to avoid blood clotting and brain drying.

#### Two-photon calcium imaging

We used a commercially available two photon microscope (A1 MP, Nikon) equipped with a 16x long-working-distance, water-immersion objective (N.A.=0.8, WD=3 mm, CFI75 LWD 16X W, Nikon) controlled by NIS-Elements software (Nikon). GCaMP6s was excited at 920 nm using a Ti:Sapphire laser system (60 fs laser pulse width; Chameleon Vision-S, Coherent). Emitted photons were collected using gated GaAsP photomultipliers (H11706-40, Hamamatsu). Movies were recorded using a resonant scanning system at a frame rate of 15 Hz and duration of 20 minutes per movie.

#### Two-photon imaging

All analysis on imaging data and treadmill behavior data were conducted in MATLAB and Python using standard toolboxes, open access toolboxes and custom written code. To remove motion artifacts, recorded movies were registered using a Lucas–Kanade model.Individual cell locations and fluorescence traces were identified using a constrained nonnegative matrix factorization based algorithm and afterwards Ca^2+^ events were identified with a constrained deconvolution algorithm 45. In order to overcome the issues of segmentation of axonal branches into smaller components an iterative merging approach was implemented. This approach was based on the correlation of the spatial footprints as well as to the temporal correlation of calcium traces of identified axons. analysesmponent extraction, those components with spatial and temporal correlations above Pearsons’ *r >* 0.2 were carefully compared individually. After inspection, components or sub-branches belonging to the same axons were merged by adding those components to each other. To follow axons over days, a 2D cross-correlation approach was utilized using standard MATLAB image registration procedures. To align the FOV between days, the astrocytic label in the red channel was used to create a rigid 2D transformation. This transformation was applied to the spatial footprints of all components. The components in the green GCaMP6s channel were then matched using the spatial footprints of each day and calculating the components with the highest 2D cross-correlation values, where maximum shift of 10 pixels was allowed between the reference and the current component. The components were assigned hierarchically and the components with *r <* 0.2 were rejected. All identified matches were inspected visually and false positive matches were rejected.

## Data analysis methods

### Data Preprocessing

Calcium image traces are smoothed using a Gaussian kernel with a size of 25 timesteps, each bin being 100ms. Timesteps where the animal is stationary are excluded in all analysis unless otherwise specified.

### Dimensionality Reduction

We used Principal Component Analysis (PCA) to reduce the dimensionality of the data. For a given session with *N*_*a*_ axons and *T* timesteps, we arrange the inferred firing rate *f*_*i*_(*t*) of axon *i* at time *t* into the activity matrix ***F*** of size (*N*_*a*_ *× T*). PCA linearly transforms the data along dimensions in descending order of explained variance

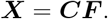

Where ***C*** is the transformation matrix and ***X*** the transformed data such that the first dimension contains the largest explained variance, the second contains the second largest, and so on. We truncated ***X*** at *N* dimensions, which we set as the amount of dimensions needed to explain a cumulative 90% of the variance (unless otherwise stated), such that ***X*** ∈ ℝ^*N ×T*^.

### Position decoding

In order to decode the position of the animal along the belt, we transformed the periodic 1-D position values to a 2-D unit circle. This allowed us to account for periodicity while having smooth, continuous position data. The angle along the unit circle corresponds to the position along the belt, using the mapping 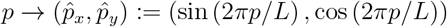 where *p* ∈ [0, 1500) is the current position in millimeters and *L* = 1500 the length of the belt. Decoding was performed using the package developed by the Kording Lab^25^, specifically the Wiener and extreme gradient boost (XGBoost) decoders. The error is taken as the standard deviation of the difference between the real and predicted positions, accounting for periodicity. In any situation where the decoder was trained and tested in the same dataset, we used 5-fold cross-validation.

### Multiset Canonical Correlation Analysis (mCCA)

In order to align the data from different sessions we used a generalization of Canonical Correlation Analysis (CCA), a classical statistical method that finds a linear transformation that maximizes the cross-correlation between two datasets. Here we adapted the algorithmic definition of a multiset CCA (mCCA) from Via et al.^26^ which uses the maximum variance (MAXVAR) definition given by Kettenring in 1971^27^, where the target is to align the maximum variance axis across data sets.

Let *M* be the number of datasets and ***X***^(*i*)^ the data matrix for dataset *i*, of size *N × T*. Our goal is to find a set of linear transformation matrices ***H***^(*i*)^ such that the sum of cross-correlations between all possible transformed dataset pairs is maximized. The space of maximum correlation between all datasets is called the “canonical” space. Let *h*^(*i*)^ be the basis vector of the ***H***^(*i*)^ transformation matrix, such that *z*^(*i*)^ = ***X***^(*i*)^*h*^(*i*)^ is the projection of dataset *i* onto the canonical dimension of maximum summed correlation. Our optimization problem reads:

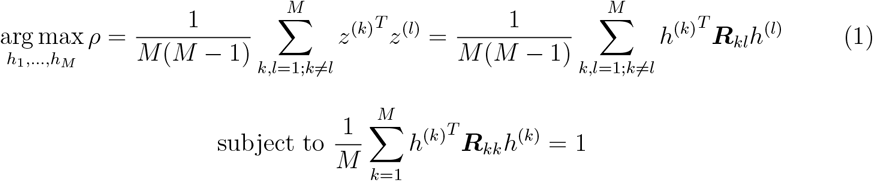

where 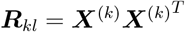 is the correlation matrix between two given datasets.

By vertically concatenating all the *h*^(*i*)^ vectors into the vector ***h*** of length *M × N*, and re-arranging terms, this optimization problem reduces to the following equation:

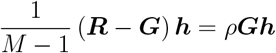

Where ***R*** is a matrix of size (*M* * *N*) *×* (*M* * *N*) containing all the *R*_*kl*_ correlation matrices, and ***G*** only the auto-correlations in its diagonal. This equation has the wellknown form of a generalized eigenvalue problem, defined as ***Av*** = *λ****Bv***, which can be solved algorithmically. In our case, 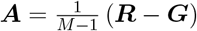, and ***B*** = ***G***.

We solve this system numerically using the *eigh* function from scikit-learn. The obtained eigenvectors *h* correspond to a concatenation of the transformation vectors *h*^(*k*)^ for each dataset. The first eigenvector transforms the data to the canonical dimension with most correlation sum, the second to the dimension of second most correlation sum, and so on. Arranging the *h*^(*k*)^ vectors from each eigenvector in rows gives us the transformation matrix ***H***^(*k*)^ for dataset *k* to the canonical space.

This mCCA solution required all datasets to be of the same size. If the number of latent dimensions in each dataset is *N* ^(*i*)^, set to explain at least 90% of the variance (details in *Dimensionality Reduction*), we truncated it at *N*_*CCA*_ = max_*i*=1,…,*M*_ *N*_*i*_ so that all datasets contained at least that much variance explained. We also needed to equalize the temporal dimension *T* ^(*i*)^, as each session has different number of trials, and each trial different number of timesteps (since the time the animal took to complete a run varied). First we warped each trial to have the same number of bins *B*, set to 150, by averaging the values within each bin. Then we truncated to the smallest number of trials *I*_*trunc*_ across sessions, so that each dataset had the common size *N*_*CCA*_ *×* (*B* * *I*_*trunc*_).

These truncated and warped arrays were only used to train the mCCA algorithm, from which we obtained the transformation matrices ***H***^(*i*)^. We quantified alignment performance using the full, unwarped datasets. There exist *M* + 1 linear spaces where the data can be aligned: the canonical space and each of the *M* starting spaces. To align to the space of a dataset *k*, we transformed all other 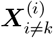 to the canonical space, and then backtransformed to the k-th space using 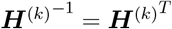. We chose the alignment space with the highest average cross-session position prediction. Each value was obtained by training a position decoder on one aligned dataset and testing on another. There are M self-predictions (tested on the training dataset) and *M* (*M* − 1) cross-predictions, done in both unaligned and aligned data. When predicting on the same session (self-prediction) 5-fold cross-validation was used.

Additionally, we added a “shifted” control that tries to align datasets that have been each temporally shifted by different random amounts, ensuring that the relationships between population activity and position are broken while internal temporal correlations remain intact. The results are averaged across 20 random combinations of shifts for each animal.

### Tensor Component Analysis (TCA)

We performed a tensor decomposition analysis using the notation of the neuron, time and trial factors introduced in Williams et al.^15^. The number of TCA dimensions was set to be the same as the number of CCA dimensions, selected to explain at least 90% of the variance on all datasets. We set the number of replicates to 10 and selected the nonnegative Parafac decomposition method with hierarchical alternative least squares(*“ncp hals”*). The optimization algorithm is not guaranteed to find a global minimum, and often multiple similar solutions are found for the same data. We repeated the analysis at least 10 times for each quantification and computed their average. Let us note that the differences for each repeat were small and did not alter the overall result.

To create the input for TCA, we truncated the aligned data at the common latent dimension and each trial normalized to have the same length (see previous section for details). Then the trials from all sessions were concatenated into a 3D tensor **𝒯** of size *N*_*CCA*_ *×I ×B*, where *I* is the total number of trials for that animal, and fed into the TCA. As a result we obtained three matrices: the dimensional factors ***D*** of size *N*_*CCA*_ *× N*_*T CA*_, the trial factors ***S*** of size *I × N*_*T CA*_, and the temporal factors ***E*** of size *B × N*_*T CA*_.

### Air puff decoding

To decode the air puff presence in a given trial, we applied the Linear Discriminant Analysis (LDA) algorithm as included in the python package `scikit-learn`^28^ on the trial factors. We chose the eigenvalue decomposition method with no shrinkage factor. Since it is a binary classification task (two labels), we projected the data onto a 1-D axis. The F1 results were computed using leave-one-out cross-validation. Additionally, to avoid data imbalance issues, multiple repeats of the same decoding were performed, choosing random subsets of the trials such that both classes always had the same number of samples. This was repeated 10 times with different trial subsets.

For the label shuffle control, class labels were randomly scrambled before training the LDA decoder, averaging the results over 10 random shuffles. We used the trial factors from the best F1 attempt in the previous step. For the “align (shift)” control, we used the data resulting from the random shifts as described in *Multiset Canonical Correlation Analysis*.

To exclude part of the belt in other control analysis, we defined an exclusion position interval and eliminated the corresponding bins from the tensor **𝒯**.

## Trial distances

For the trial distance analysis, we computed Euclidean distances between each trial factor, corresponding to the rows in matrix ***S***, and the center of mass of each class (defined by either session type, or the presence of an air puff). For these calculations, we set *N*_*TCA*_ to be 11 for all mice, as to compare distances in the same N-dimensional space. We chose this value as the average across mice to explain 90% of the variance.

For the within-cluster distance significances, we first calculated the difference between either B or T distances to P, and then did significance testing on the ID group difference against the DD group difference.

### Decoding weights

To quantify the contribution of a feature (axon or latent dimension) to a given decoding, we used the weights from linear decoders for position and air puff decoding.

### Position weights

For position, we used the coefficients obtained from Wiener filter regression, which for the predicted position 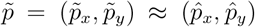 are given by the weights *ω* in:

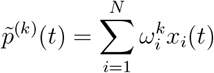

where *k* = *x* or *y* (one of the two coordinates after the 2D transformation), *x*_*i*_(*t*) is the value of feature *i* at time *t* (in our case, the PCA dimensions), and *N* is the total number of features considered in the analysis (see *Position Decoding* for details). We took the contribution of PCA dimension *i* to position decoding as the average 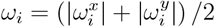

Given the PCA transformation matrix ***C*** (see *Dimensionality Reduction*), its elements *c*_*ij*_ denote the contribution of cell *j* = 1, …, *N*_*a*_ to principal component *i*. And conversely, since ***C*** is orthogonal, *c*_*ji*_ is the loading of dimension *i* onto cell *j*, such that its reconstructed firing rate is given by *f*_*j*_(*t*) = _*i*_ *c*_*ji*_*x*_*i*_(*t*). We then defined the contribution of cell *j* to position decoding as:

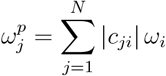

Intuitively, if |*c*_*ji*_| is relatively large, cell *j* is a major contributor to variations in principal component *i*, and hence a main contributor to decoding weight *ω*_*i*_.

Finally, weights 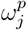 were z-scored to allow comparison between different sessions and animals.

### Air puff weights

Air puff presence was decoded using LDA (see *Air puff decoding*). Its input are the TCA trial factors corresponding to the rows *S*_*i*_ in matrix ***S*** (see *Tensor Component Analysis (TCA)*), each of length *N*_*T CA*_. Since our classification problem is binary, LDA projects the data to a 1-D space using the vectors *ω*^*LDA*^:

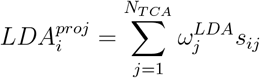

Where 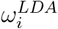 is the LDA weight for TCA dimension *j* and *s*_*ij*_ the element *ij* from ***S***. The weights were averaged over cross-validation folds and data balancing repetitions, to obtain a final *ω*^*LDA*^ weight vector.

Next, we computed the decoding contribution of each CCA aligned dimension using the dimensional factors ***D***, each element *d*_*ij*_ being the loading of CCA dimension *i* to TCA dimension *j*, with the formula:

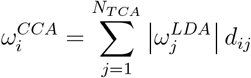

Obtained weights were averaged over TCA repetitions.

Having the contribution of each CCA dimension, we used the transformation matrices ***H***^(*k*)^ to estimate the contribution of each PCA dimension *i* from session *k* to air puff decoding with the formula:

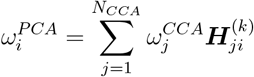

Where we used the backtransformation matrices 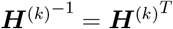

Then, to obtain the contribution of each cell to air puff decoding, we back-transformed from PCA space to axonal space, again using the inverse ***C***^*−*1^ = ***C***^*T*^ :

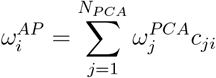

To summarize, the contribution to air puff decoding can be reduced to the following matrix multiplication:

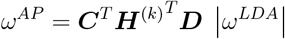

Where | · | represents applying the absolute operator element-wise. As with the position weights, the air puff weights 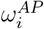 were z-scored to allow comparison between different sessions and animals.

Place and non-place cell classification was done using the Dombeck standard^29^.

## Supplementary Figures

**Figure S1:**
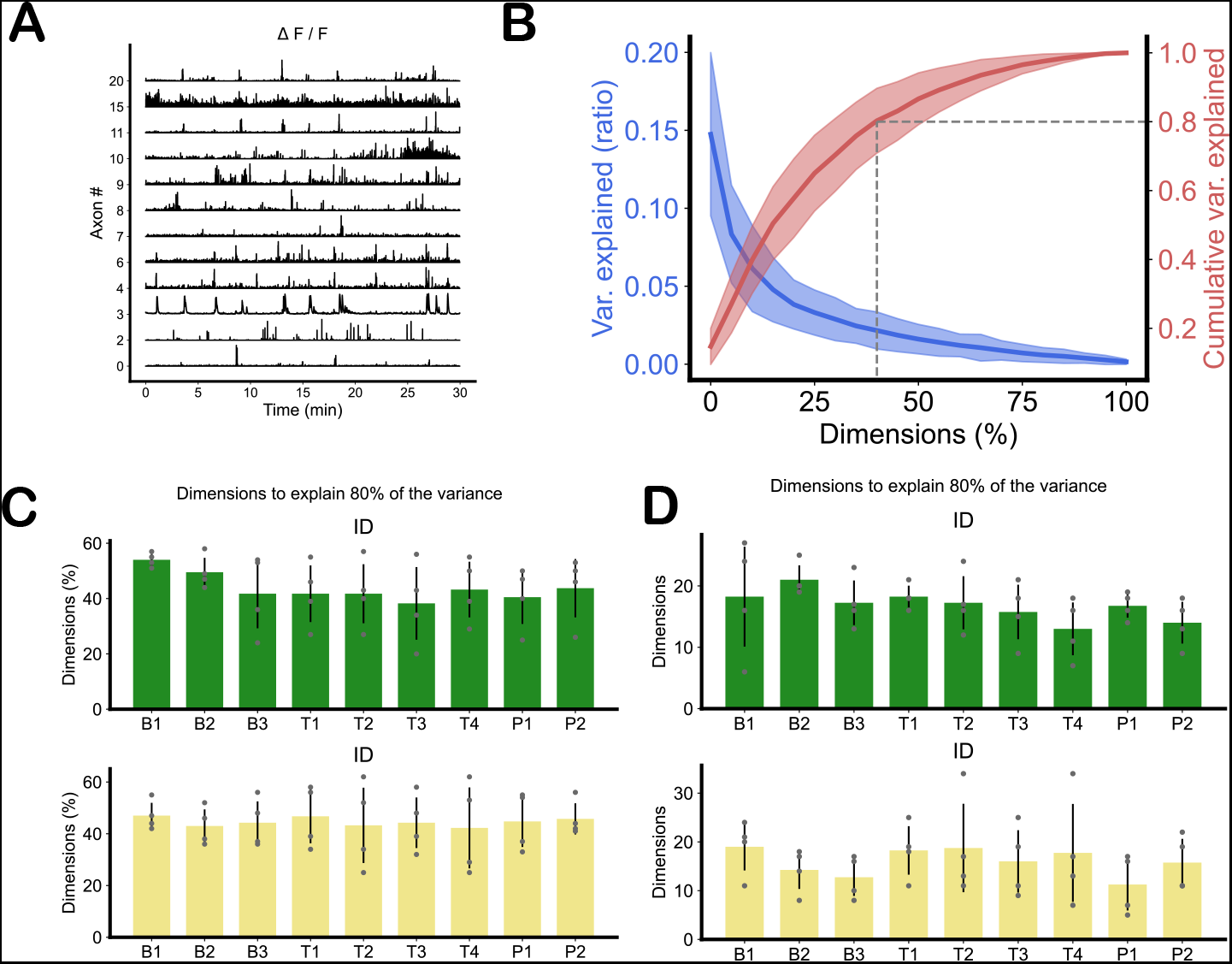
Raw data and dimensionality plots. **A**, Example of a Δ*F/F* fluorescent signal recorded during a session. **B**, Variance ratio and cumulative variance explained by dimension, averaged over all mice and sessions. Shaded area represents the standard deviation of the data. Dashed line shows the average percentage of dimensions needed to explain 80% of the variance. **C**, Average percentage of dimensions needed to explain 80% of the variance for each session and axon type. Error bar represents the standard deviation. Results are averaged over 4 mice for each, shown as gray dots. Overall we needed (44% ± 10%(SD)) to explain 80% of the variance. **D**, Same as in **C** but considering the absolute number of dimensions instead of a given percentage.

**Figure S2:**
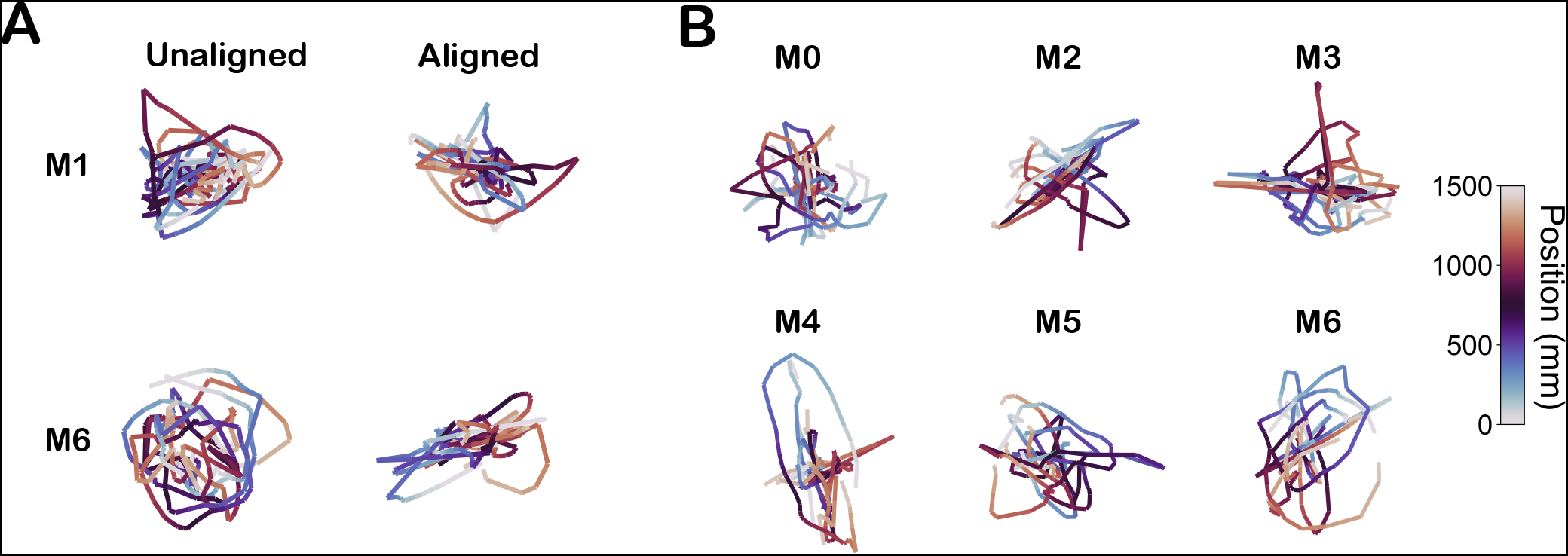
Alignment controls. **A**, Each row shows the trial-average PCA of each session before and after alignment for “shifted” recordings. Each session has been shifted by a random amount, and then aligned using CCA. Since neuronal activity has a different encoding with position for every session, the alignment process fails. This is shown for two mice (M1 and M6) and one random selection of session shifts for each mouse. **B**, Trial-averaged PCAs of each session after alignment for “shifted” recordings, for each remaining mice. As in the previous panel, this shows a single random selection of session shifts for each mouse.

**Figure S3:**
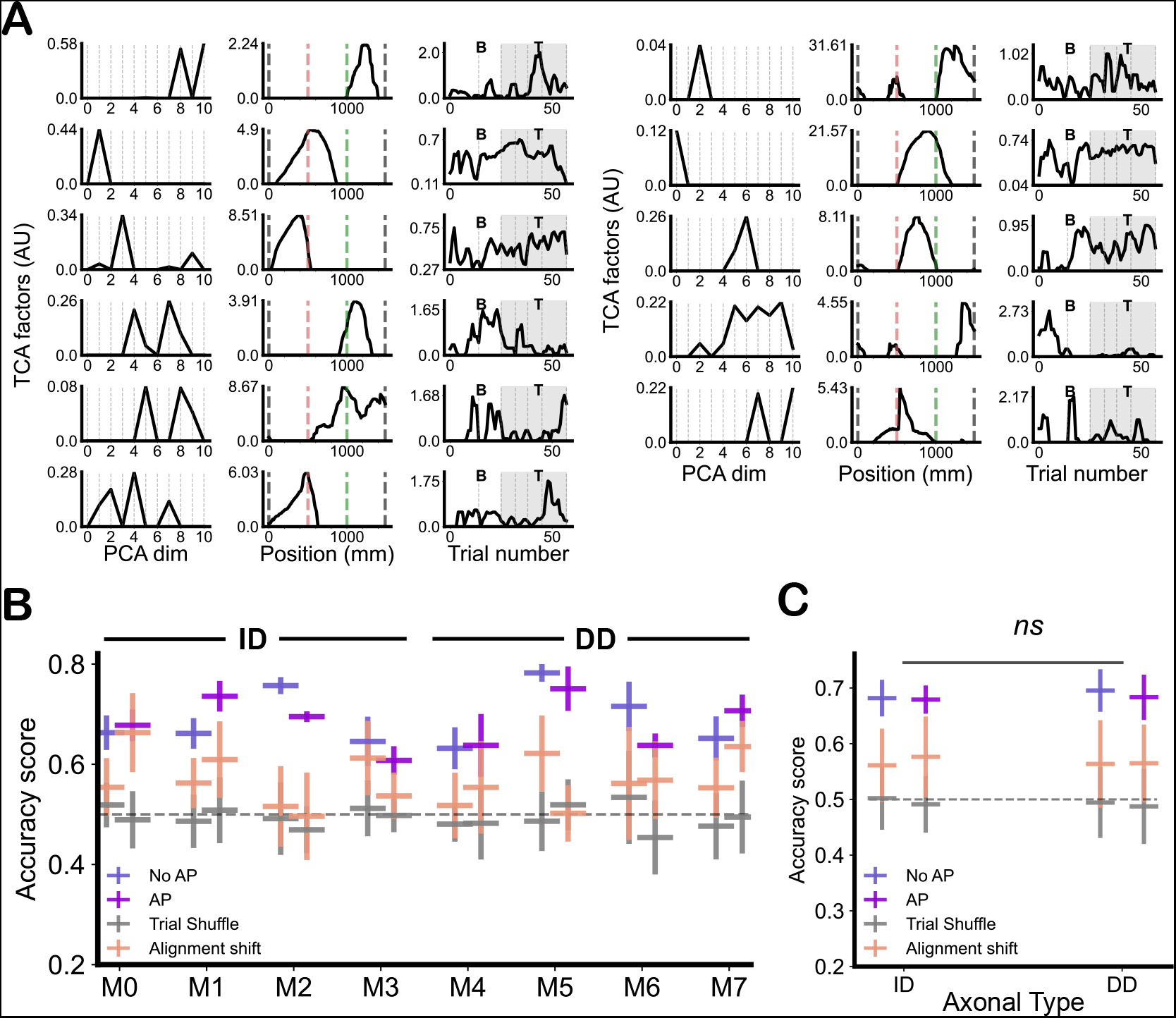
Air puff decoding accuracy. **A**, TCA factors for mouse 6 (DD). As many factors as dimensions needed to explain at least 90% of the variance on each session were chosen, in this case *N*_*T CA*_ = 11. Left column shows the PCA factors. middle column positions factors with dashed lines indicating the reward (gray), air puff (red), and last belt transition (green) locations. Third column shows the trial factors for baseline (B) and training (T) trials. Trials where there air puff is present are colored in gray. **B**, Mouse-averaged raw accuracy scores (proportion of correct labels) for each mouse and class. Dashed line represents the expected chance of random guessing. Vertical bars represent the standard deviation of the data. **C**, Axonal-type-averaged raw accuracy scores (proportion of correct labels) for each mouse type and class. Dashed line represents the expected chance of random guessing. Vertical bars represent the standard deviation of the data.

**Figure S4:**
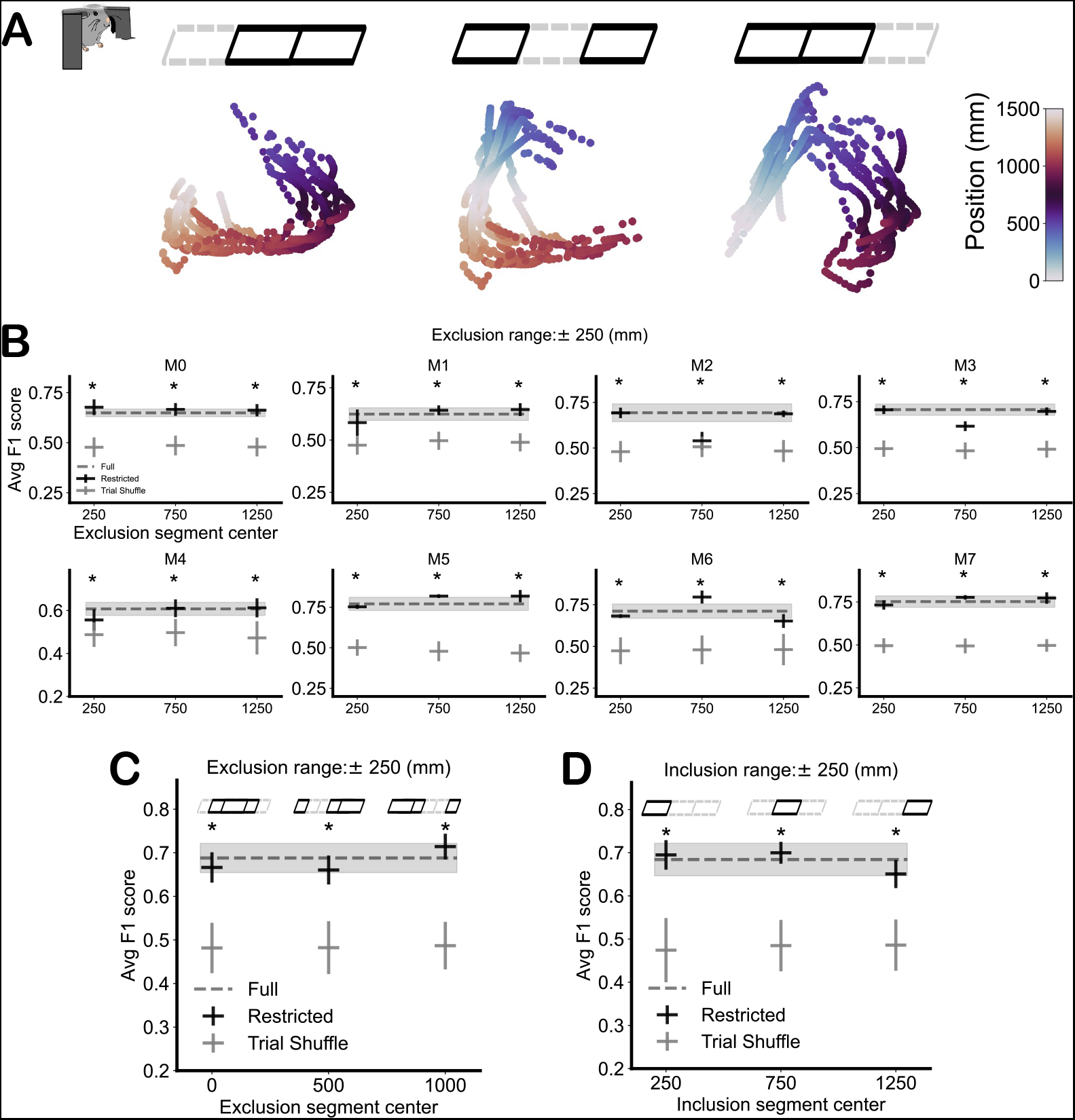
Excluding sections of the belt does not change air puff decodability results. **A**, Example PCAs from mouse 6 after excluding one of three sections fo the periodic belt, colored according to position. The top drawings show which third of the belt was excluded. **B**, F1 scores after excluding one third of the belt for each mouse. The dashed line and the gray area show the average F1 scores and their standard deviation, respectively, when no exclusion is performed. Asterisks represent significance with shuffle (t-test, *p <* 0.05). **C**, Mouse-averaged F1 scores after excluding one third of the belt, but centered around belt section transitions. There was no significant drop in decodability, indicating that the overall results were not affected by the stopping behavior of the animal. **D**, Mouse-averaged F1 scores after excluding two thirds of the belt. Once again there was no significant drop in decodability.

**Figure S5:**
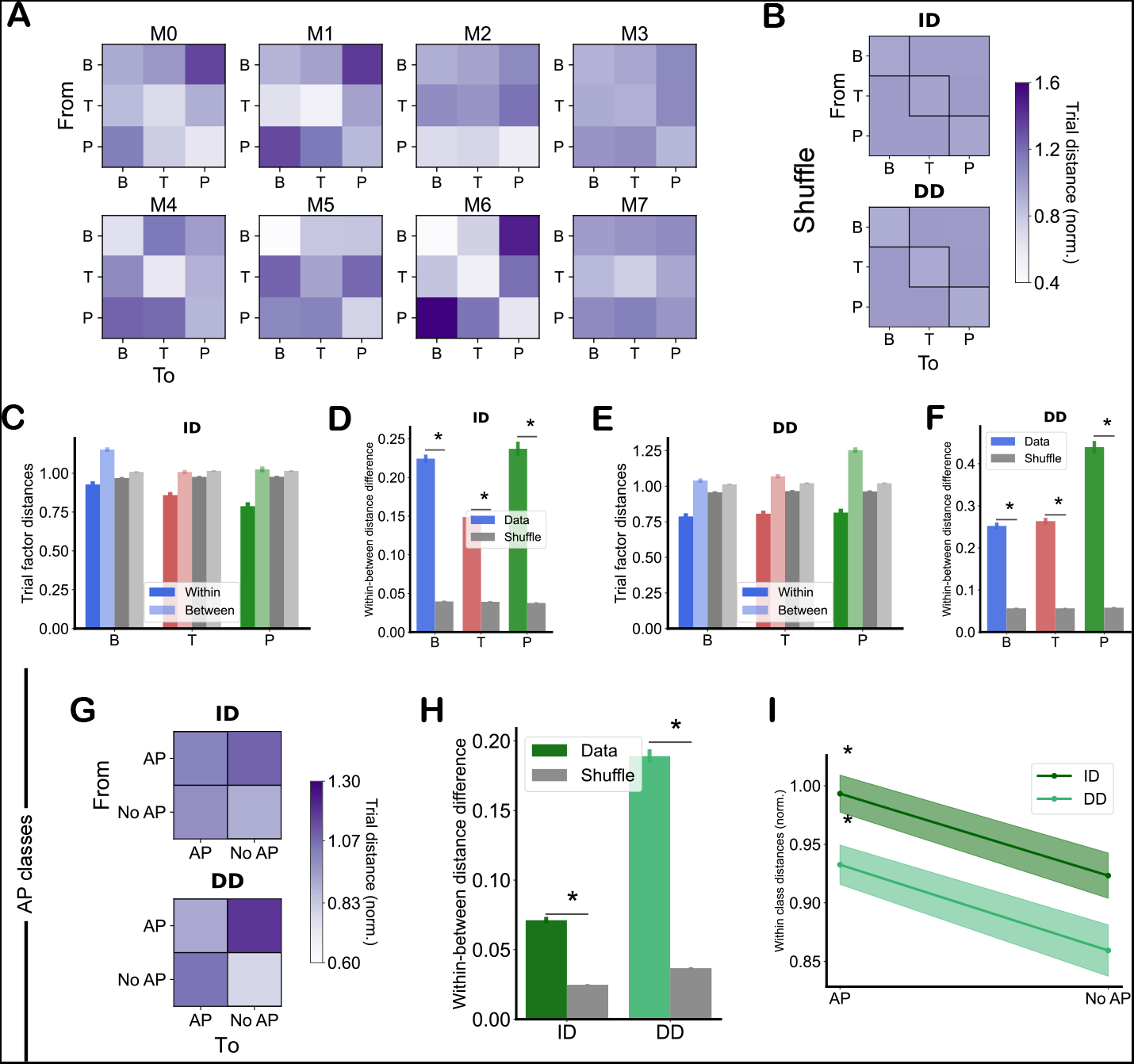
Euclidean distance calculations for trial factors. **A**, Colormaps of the average normalized distance from a trial to each centroid, by mouse. The diagonal elements show the average within-cluster distance. **B**, Colormaps of the average normalized distance from a trial to each centroid, by axonal type, after random trial label shuffling. The diagonal elements show the average within-cluster distance. This colormap shows more homogeneity than panel A or Fig. 4C. **C**, Mouse-normalized average distances between a trial factor and the center of mass of its own session type (“within”) or the center of masses of the other session types (“between”), for each session type. Gray bars represent the shuffled results. Vertical lines show the standard error of the mean. Results shown for ID axons only. **D**, The “within” and “between” distances from the previous panel are subtracted in order to quantify cluster separability. This panel shows the mouse-normalized average distance differences for each session type. Gray bars represent the shuffled results. Asterisks indicate significant (t-test, *p <* 0.05). Here, results shown for ID axons only. **E**,**F** Same as panels C,D but for DD axons. **G** Same as panel A, but this time grouping trials by air puff presence (T sessions, “AP”) and no air puff presence (B and P sessions, “NoAP”). **H** Average difference between the “within” and “between” normalized distances from panel G, for each axonal type, for both the original data and the label-shuffle. Shaded areas represent the standard error of the data. Significance (t-test, *p <* 0.05) was computed with respect to label-shuffle. **I** Average distances within a cluster for AP and NoAP classes. Asterisk indicates significance (t-test, *p <* 0.05) between the two classes.

**Figure S6:**
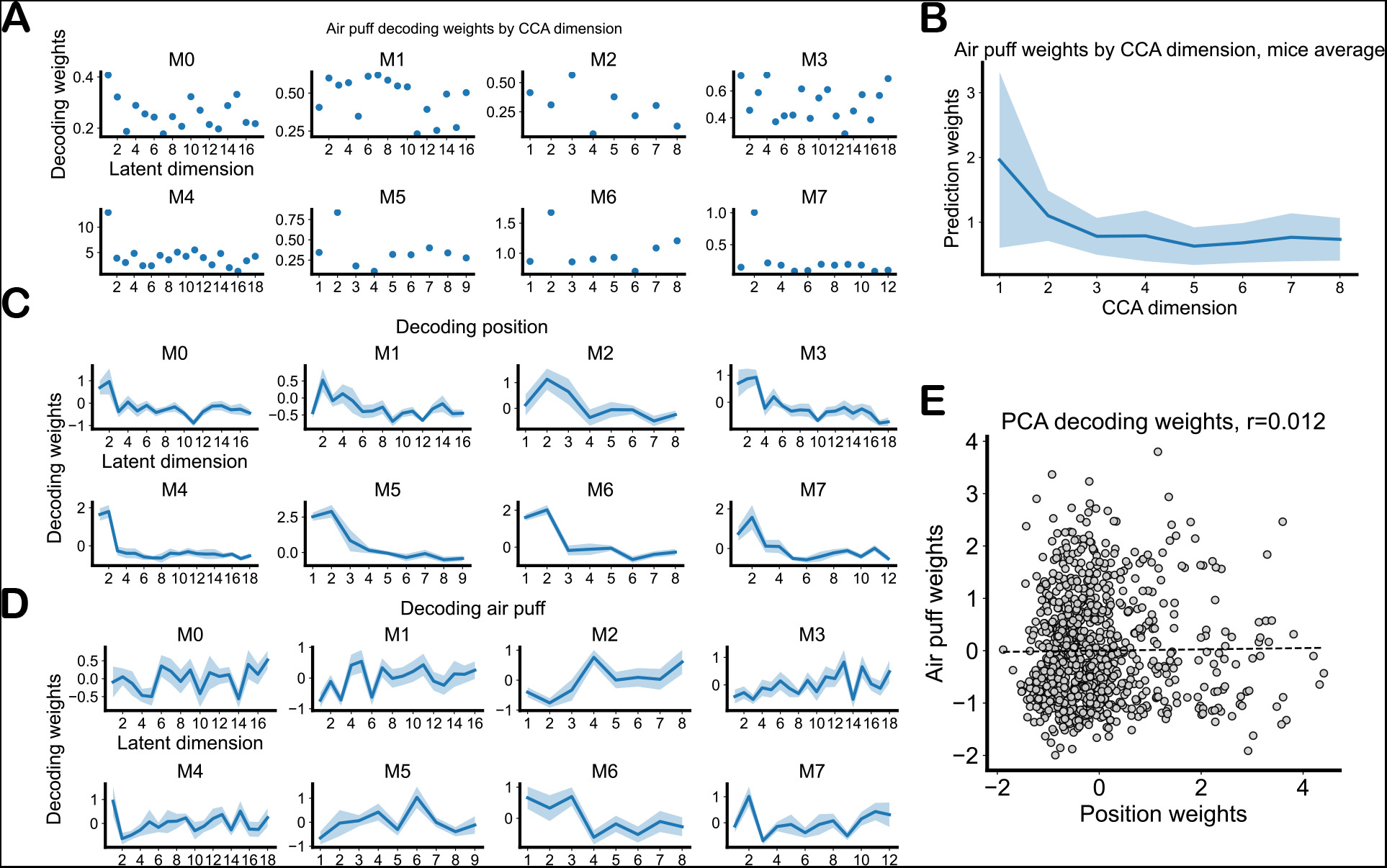
Decoding weights for position and air puff. **A**, Air puff decoding weights for each aligned CCA dimension and mouse. **B**, Average normalized air puff decoding weights by CCA latent dimension across mice. Shaded area represents the standard error of the mean. **C**, Average, session-normalized position decoding weights by PCA dimension for each mouse. Shaded area represents the standard error of the mean. **D**, Same as C, but for air puff decoding weights. **E** Plotting air puff decoding weights against position decoding weights for each PCA dimension. Dashed line shows linear regression, which was non-significant with a p-value of 0.28.

