## Supplementary Figures for "Transformations of the spatial activity manifold convey aversive information in CA3"

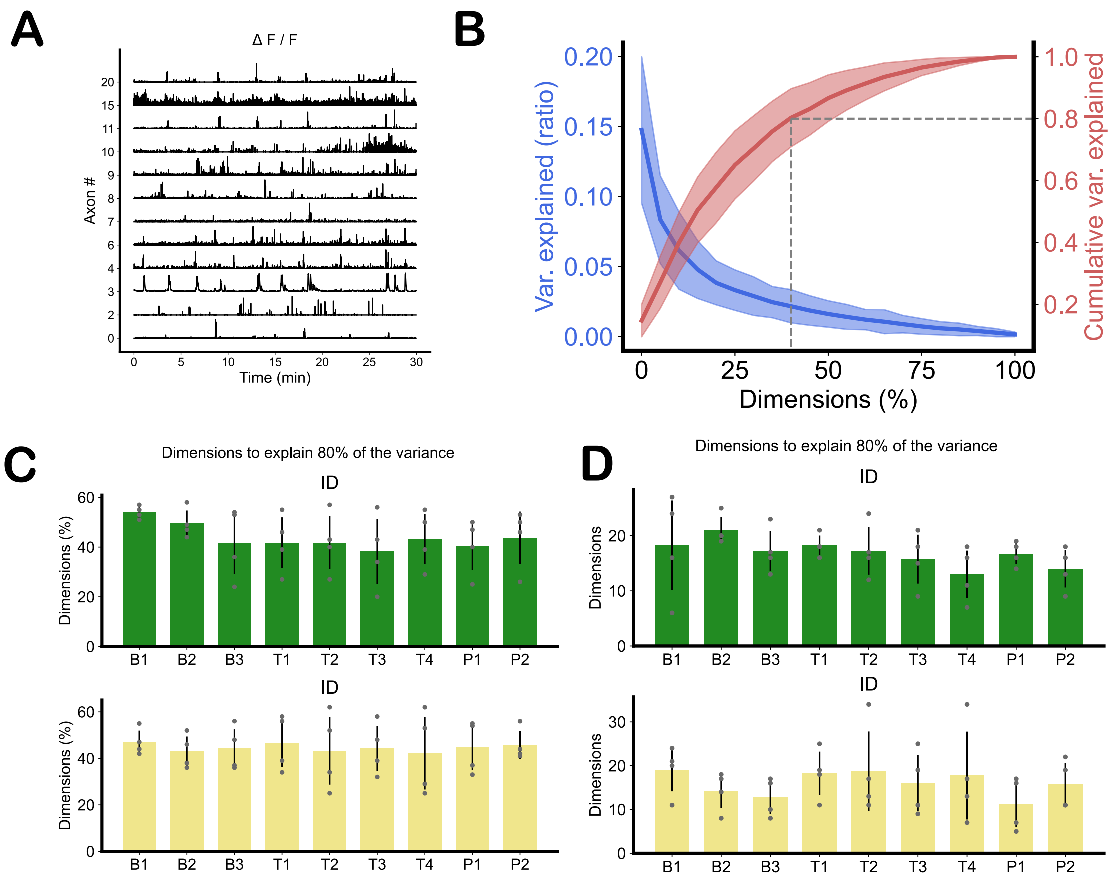


Figure S1: **Raw data and dimensionality plots. A,** Example of a $\Delta F/F$ fluorescent signal recorded during a session. **B,** Variance ratio and cumulative variance explained by dimension, averaged over all mice and sessions. Shaded area represents the standard deviation of the data. Dashed line shows the average percentage of dimensions needed to explain 80% of the variance. **C,** Average percentage of dimensions needed to explain 80% of the variance for each session and axon type. Error bar represents the standard deviation. Results are averaged over 4 mice for each, shown as gray dots. Overall we needed ($44\% \pm 10\%$(SD)) to explain 80% of the variance. **D,** Same as in **C** but considering the absolute number of dimensions instead of a given percentage.

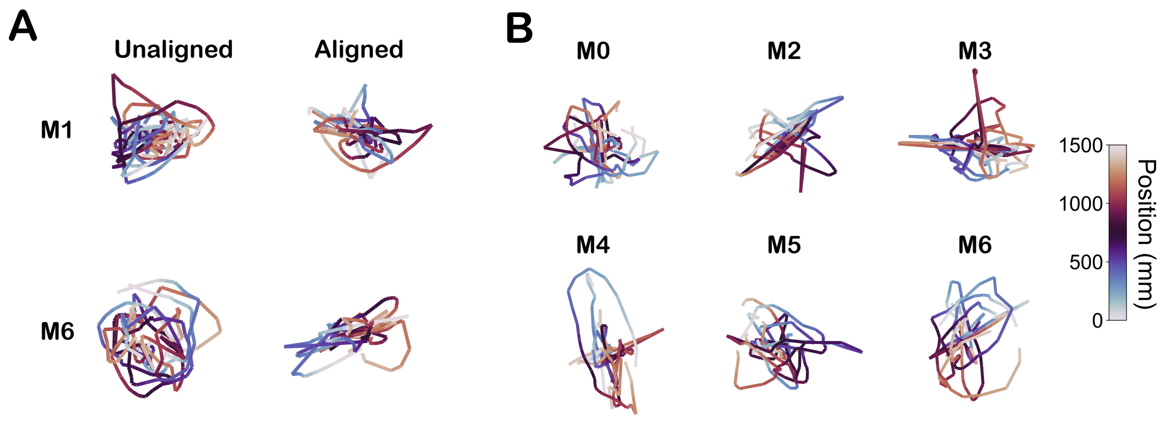


Figure S2: **Alignment controls.** **A,** Each row shows the trial-average PCA of each session before and after alignment for "shifted" recordings. Each session has been shifted by a random amount, and then aligned using CCA. Since neuronal activity has a different encoding with position for every session, the alignment process fails. This is shown for two mice (M1 and M6) and one random selection of session shifts for each mouse. **B,** Trial-averaged PCAs of each session after alignment for "shifted" recordings, for each remaining mice. As in the previous panel, this shows a single random selection of session shifts for each mouse.

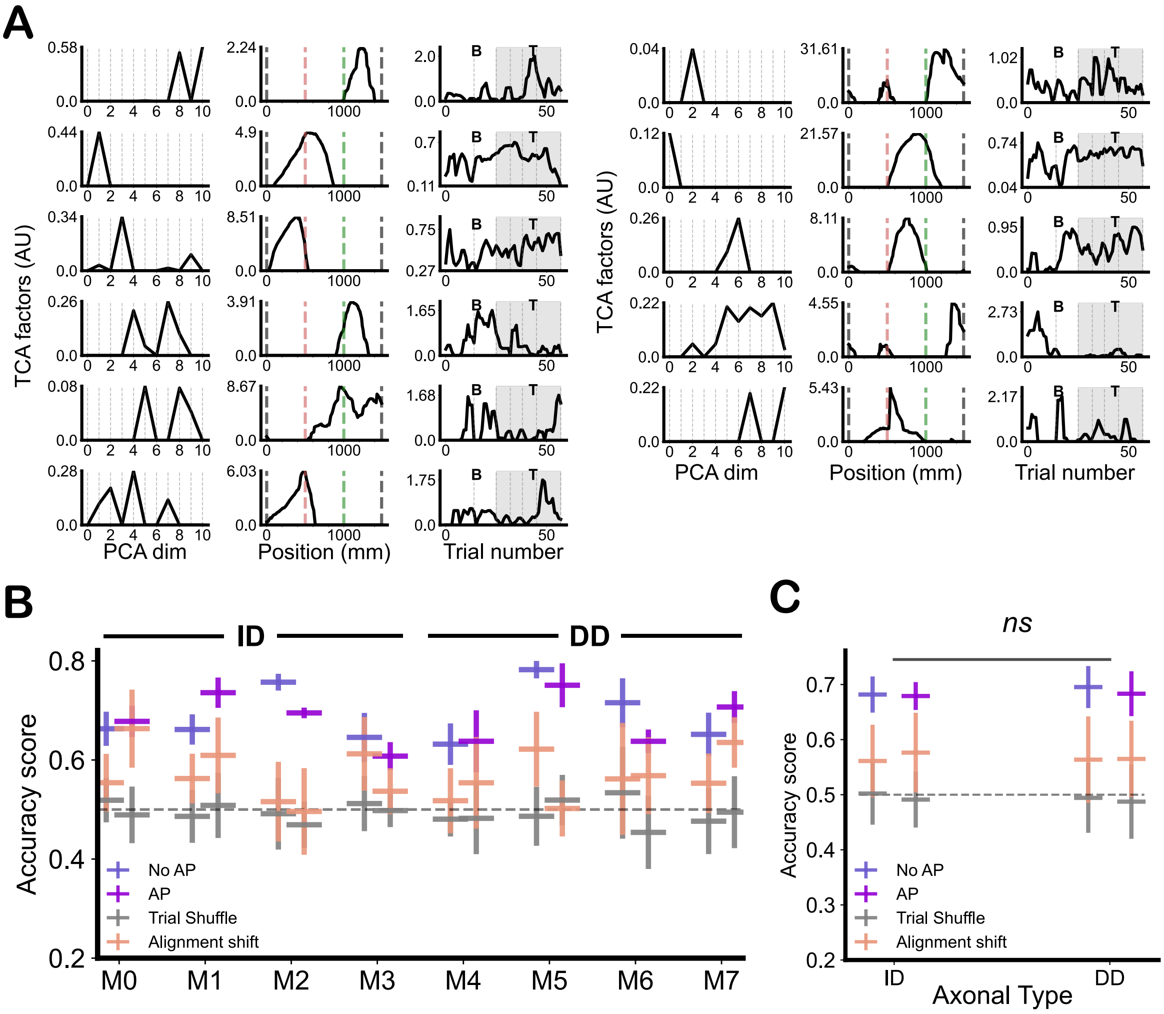


Figure S3: **Air puff decoding accuracy. A,** TCA factors for mouse 6 (DD). As many factors as dimensions needed to explain at least 90% of the variance on each session were chosen, in this case $N_{TCA} = 11$. Left column shows the PCA factors. middle column positions factors with dashed lines indicating the reward (gray), air puff (red), and last belt transition (green) locations. Third column shows the trial factors for baseline (B) and training (T) trials. Trials where there air puff is present are colored in gray. **B,** Mouse-averaged raw accuracy scores (proportion of correct labels) for each mouse and class. Dashed line represents the expected chance of random guessing. Vertical bars represent the standard deviation of the data. **C,** Axonal-type-averaged raw accuracy scores (proportion of correct labels) for each mouse type and class. Dashed line represents the expected chance of random guessing. Vertical bars represent the standard deviation of the data.

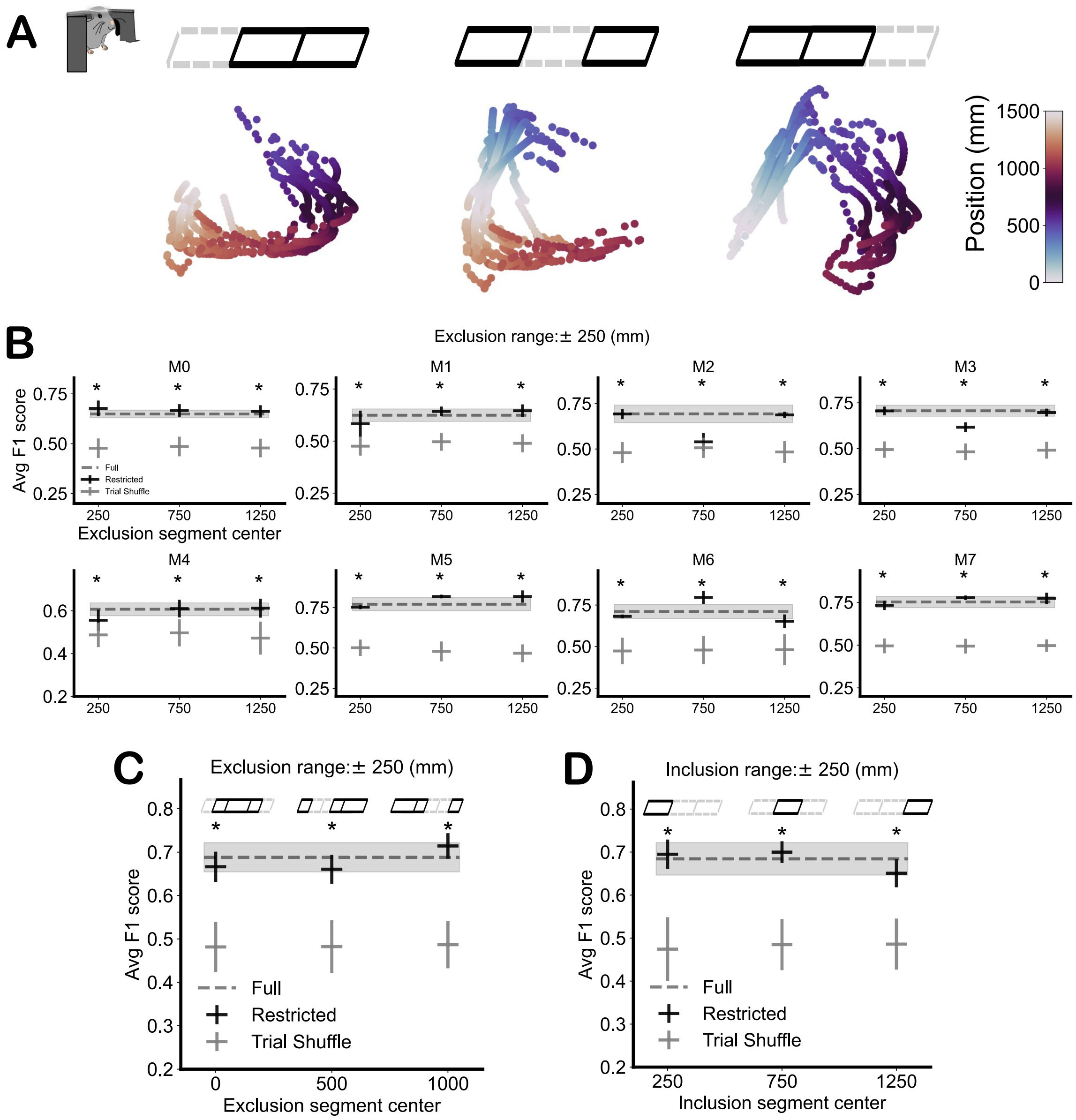


Figure S4: **Excluding sections of the belt does not change air puff decodability results.** **A,** Example PCAs from mouse 6 after excluding one of three sections fo the periodic belt, colored according to position. The top drawings show which third of the belt was excluded. **B,** F1 scores after excluding one third of the belt for each mouse. The dashed line and the gray area show the average F1 scores and their standard deviation, respectively, when no exclusion is performed. Asterisks represent significance with shuffle (t-test, $p < 0.05$). **C,** Mouse-averaged F1 scores after excluding one third of the belt, but centered around belt section transitions. There was no significant drop in decodability, indicating that the overall results were not affected by the stopping behavior of the animal. **D,** Mouse-averaged F1 scores after excluding two thirds of the belt. Once again there was no significant drop in decodability.

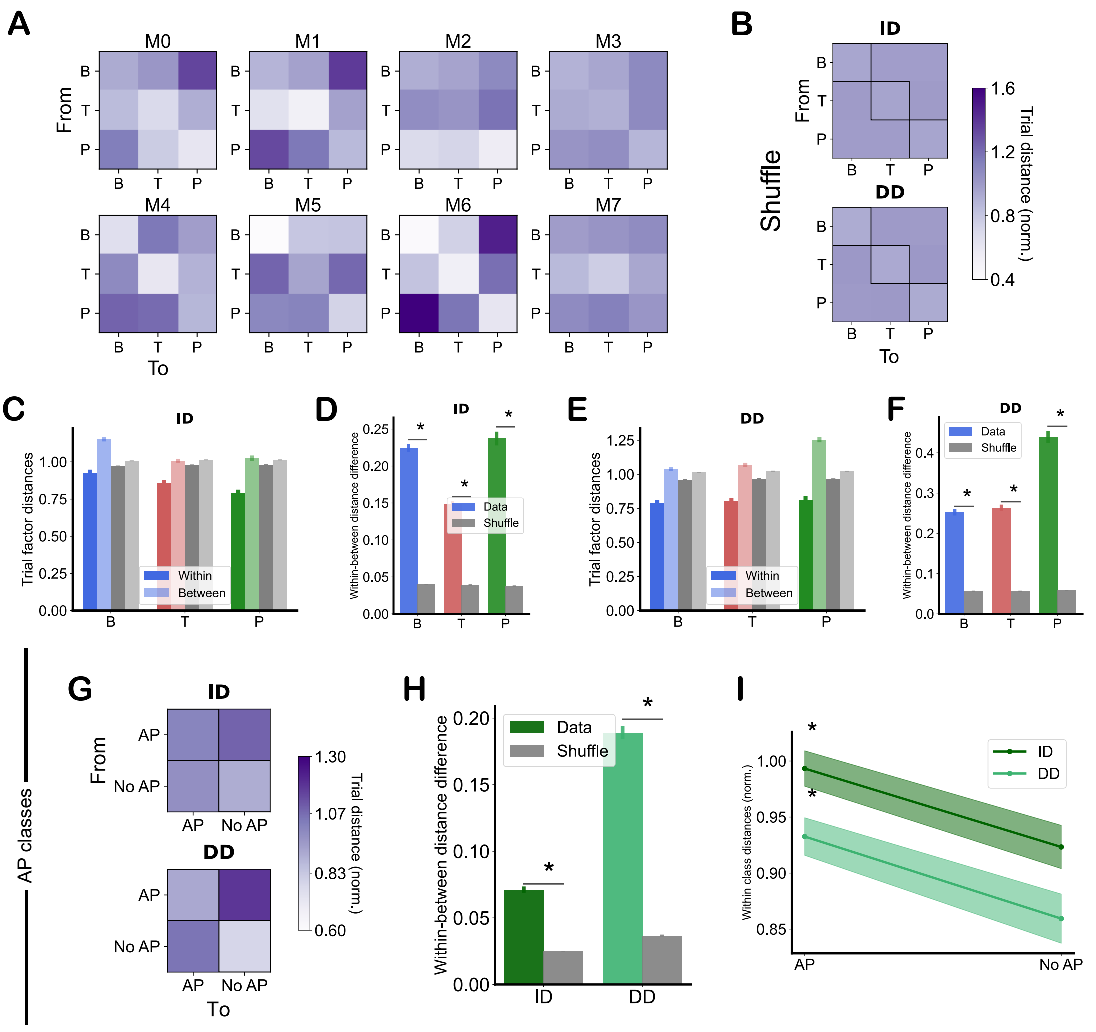

Figure S5: **Euclidean distance calculations for trial factors. A,** Colormaps of the average normalized distance from a trial to each centroid, by mouse. The diagonal elements show the average within-cluster distance. **B,** Colormaps of the average normalized distance from a trial to each centroid, by axonal type, after random trial label shuffling. The diagonal elements show the average within-cluster distance. This colormap shows more homogeneity than panel A or Fig. 4C. **C,** Mouse-normalized average distances between a trial factor and the center of mass of its own session type ("within") or the center of masses of the other session types ("between"), for each session type. Gray bars represent the shuffled results. Vertical lines show the standard error of the mean. Results shown for ID axons only. **D,** The "within" and "between" distances from the previous panel are subtracted in order to quantify cluster separability. This panel shows the mouse-normalized average distance differences for each session type. Gray bars represent the shuffled results. Asterisks indicate significant (t-test, $p < 0.05$). Here, results shown for ID axons only. **E,F** Same as panels C,D but for DD axons. **G** Same as panel A, but this time grouping trials by air puff presence (T sessions, "AP") and no air puff presence (B and P sessions, "NoAP"). **H** Average difference between the "within" and "between" normalized distances from panel G, for each axonal type, for both the original data and the label-shuffle. Shaded areas represent the standard error of the data. Significance (t-test, $p < 0.05$) was computed with respect to label-shuffle. **I** Average distances within a cluster for AP and NoAP classes. Asterisk indicates significance (t-test, $p < 0.05$) between the two classes.

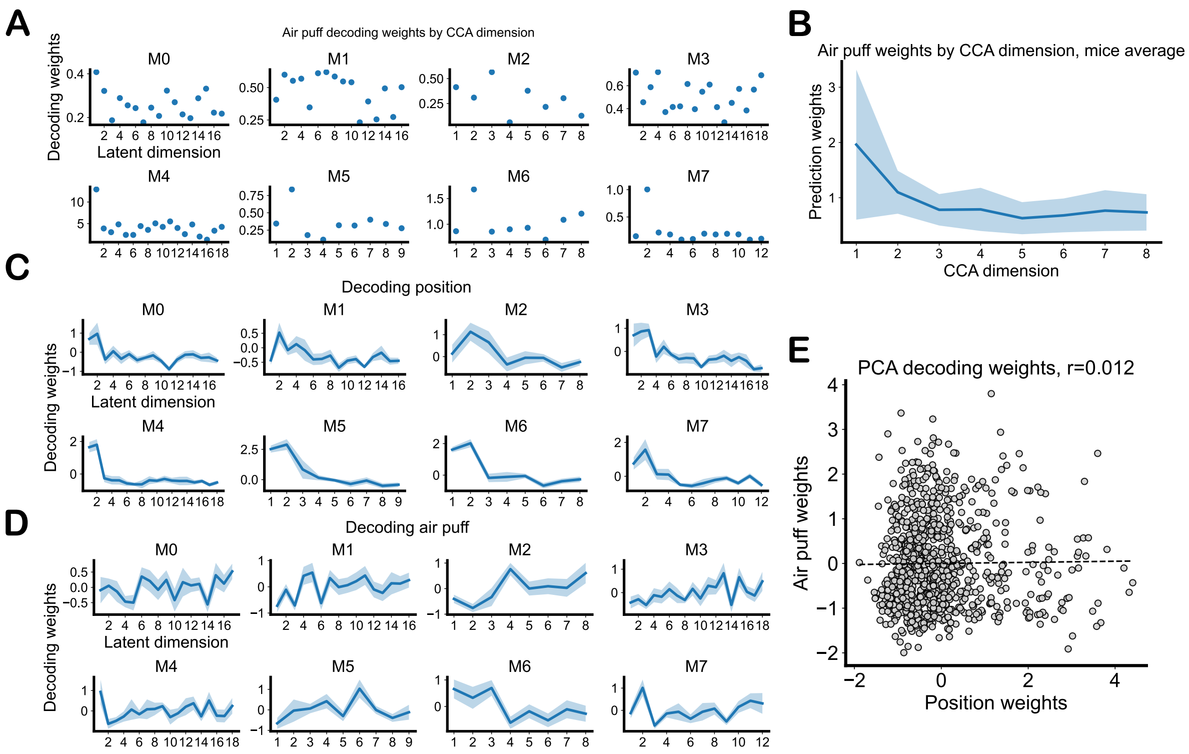


Figure S6: **Decoding weights for position and air puff. A,** Air puff decoding weights for each aligned CCA dimension and mouse. **B,** Average normalized air puff decoding weights by CCA latent dimension across mice. Shaded area represents the standard error of the mean. **C,** Average, session-normalized position decoding weights by PCA dimension for each mouse. Shaded area represents the standard error of the mean. **D,** Same as C, but for air puff decoding weights. **E** Plotting air puff decoding weights against position decoding weights for each PCA dimension. Dashed line shows linear regression, which was non-significant with a p-value of 0.28.
